# Two-brain states characterise within- and between-brain connectivity during intergenerational collaborative drawing

**DOI:** 10.64898/2026.01.29.702550

**Authors:** Luca A. Naudszus, Ryssa Moffat, Emily S. Cross

**Affiliations:** Social Brain Sciences Lab, ETH Zurich

**Author notes:** **Corresponding Authors:** Luca A. Naudszus, Emily S. Cross, **Address:** Social Brain Sciences Lab, ETH Zurich, Stampfenbachstrasse 69, Zurich 8092, Switzerland. Equal contribution (Luca A. Naudszus, Ryssa Moffat). **Author Contributions** LAN: Conceptualisation, Data curation, Formal analysis, Investigation, Validation, Visualisation, Writing – original draft, Writing – review & editing. RM: Supervision, Equipment donation, Conceptualisation, Data curation, Investigation, Visualisation, Writing – original draft, Writing – review & editing. ESC: Conceptualisation, Funding acquisition, Project administration, Writing – review & editing.

**Keywords:** fNIRS, intergenerational, creative drawing, functional connectivity, brain states, collaboration, dyadic

## Abstract

Engaging with others in social scenarios can result in the alignment of brain activity between individuals. Dyadic (i.e., hyperscanning) studies typically estimate average region-specific levels of connectivity between brains (as opposed to within brains) across a given task to quantify brain activity alignment. This approach assumes symmetric interactions with equal and mutual adaptation from both dyad members, excluding asymmetric (e.g., leader–follower) contexts. Such approaches also obscure spatial dynamics (i.e., relationship between different brain regions) and temporal dynamics during unfolding interactions. To overcome these challenges, we took a data-driven approach to quantify within- and between-brain connectivity during collaborative drawing among dyads. Specifically, we used sliding windows and Riemannian-geometry-based *k*-means clustering to identify recurrent two-brain states while 61 dyads drew alone and together at 6 weekly timepoints. Thirty dyads comprised young adults only (same generation) and 31 dyads comprised one older and one younger individual (intergenerational). We identified 7 two-brain states, 3 of which were specific to real (not pseudo) dyads. One two-brain state showed convincing evidence of sensitivity to collaboration context: During collaborative drawing, low-to-medium between-brain connectivity and prominent within-brain connectivity in bilateral IFG in a single dyad member arose for longer periods in intergenerational than same generation dyads (co-occurring with reduced turn-taking behaviour). No two-brain state showed evidence of longitudinal changes across sessions. These findings inform recent accounts of neural dynamics that emphasise the complementary roles of within-brain and between-brain connectivity. Furthermore, they suggest that state-based analyses can inform neural dynamics in a way not captured by traditional analysis techniques.

**Highlights:**

- Two-brain states can characterise within- and between-brain connectivity.
- Seven two-brain states identified using data-driven approach.
- Intergenerational collaborative drawing linked to asymmetric connectivity.

## 1 Introduction

Human social interaction is an inherently dynamic and reciprocal process, whether between people of similar or very different ages. Advances in wearable brain imaging grant increasingly sophisticated insights into the neural underpinnings of rich, real-world interactions between dyads and groups of people. To do so, studies typically quantify levels of interpersonal neural synchrony, by estimating the level of connectivity between interacting individuals’ brains. This approach is primarily suited to understand symmetric interactions where both dyad members display the same underlying processes (e.g., mutual adaptation). However, social interactions are often characterised by high asymmetry (e.g., in leader–follower dynamics) and switching roles. To further understand the neural dynamics of social interactions, methods that integrate within-brain and between-brain connectivity while preserving temporal information are needed. In this study, we apply a state-based approach to functional near-infrared spectroscopy (fNIRS) data, recorded while intergenerational and same generation dyads completed collaborative drawing tasks, to demonstrate the utility of this method. In the following sections, we briefly describe interpersonal neural synchrony, current theoretical and empirical stances on within-brain and between-brain connectivity, recent advances in state-based analyses for dyadic neural data, before introducing the present study.

### 1.1 Interpersonal alignment and interpersonal neural synchrony

During social interactions, various forms of interpersonal alignment can arise between the interacting people (De Felice et al., 2025; Hasson et al., 2012; Schilbach et al., 2013; Shamay-Tsoory et al., 2019). Interpersonal neural alignment–the focus of this work–refers to the temporal synchronisation of patterns of brain activity between individuals. Interpersonal alignment can also occur at the level of cognition (e.g., emotional contagion, alignment in perceptions or beliefs), motor behaviours (e.g., matching body postures or movements), physiological signals (e.g., heart rate or electrodermal activity), and endocrine (e.g. oxytocin) trajectories (Shamay-Tsoory et al., 2019). Interpersonal alignment is theorised to be an adaptive mechanism that supports human health and wellbeing by maintaining social cohesion (Bilek et al., 2022; Dikker et al., 2019, 2021; Hoehl et al., 2021).

Interpersonal neural alignment can be quantified using measures of interpersonal neural synchrony (INS), derived from simultaneously recorded brain signals belonging to two or more interacting people. The acquisition of such simultaneous recordings is commonly referred to as hyperscanning. Though hyperscanning can technically be conducted with any brain imaging technique, mobile techniques such as functional near-infrared spectroscopy (fNIRS), electroencephalography (EEG) are commonly used, as their mobility is optimal for naturalistic social interactions involving body movement (Czeszumski et al., 2020; Hakim et al., 2023).

Collaboration relies on people dynamically tracking how their own behaviour influences their collaborative partner’s behaviour (Czeszumski et al., 2022; Happé et al., 2017; Schurz et al., 2021). Cortical regions implicated in processing social information and collaboration include the inferior frontal gyrus (IFG) and temporoparietal junction (TPJ; Czeszumski et al., 2022). The IFG is a key node of the action observation network (Cross et al., 2009; Shamay-Tsoory, 2022), which shows enhanced INS during joint attention (Redcay et al., 2012; Yoshioka et al., 2021) and interpersonal coordination (Marton-Alper et al., 2023). The TPJ is a core node of the Theory of Mind network, underpinning perspective taking and integration of social information about oneself and other, both core components of social cognition (Jacoby et al., 2016; Saxe & Kanwisher, 2003). Prior work has shown both of these processes to be reflected in INS levels during collaboration (Bilek et al., 2015; Czeszumski et al., 2022; Goelman et al., 2019). To understand core components of social cognitions, such as joint attention, perspective taking, and integration of social information can shape dynamic collaborative social interactions, hyperscanning studies have dominantly recorded brain signals from the IFG and TPJ.

### 1.2 Theoretical and empirical approaches to within-brain and between-brain connectivity

Two main theoretical frameworks account for the emergence of INS, respectively termed common cognitive processing and mutual prediction. *Common cognitive processing* posits that when two people interact, both people perceive similar stimuli, causing both people’s brains to respond in comparable ways at comparable time delays (Burgess, 2013). This framework is limited to characterising symmetrical forms of interpersonal alignment, such as shared attention or mutual coordination (Hamilton, 2021). Social interactions may, however, take many additional forms (e.g., individual action where one person demonstrates something for another person to learn, asymmetric adaptation, and turn-taking). Asymmetric social interactions may arise from different physical roles, such as speaking and listening, teaching and learning, demonstrating and imitating, or different social roles, such as pairings of clinicians and patients, parents and children, or even people of different generations or cultures. Of note, a wealth of hyperscanning studies examine symmetrical and asymmetrical interactions between parents and children, as well as students and teachers (DePasquale, 2020; L. Zhang et al., 2022). While such studies shed light on the neural dynamics of social interactions of common intergenerational pairs (i.e., parent–child and teacher– child), the neural dynamics of intergenerational interactions involving older adults have yet to receive a similar degree of empirical attention, despite older adults representing a large proportion of today’s global population (Moffat, 2024; Pauly et al., 2021; United Nations, 2024). Moreover, a deeper understanding how older adults’ brains negotiate collaborative social interactions will contribute to a comprehensive understanding of the social brain in later life.

Relative to common cognitive processing, the *mutual prediction* framework is adapted for symmetric and asymmetrical interactions. The mutual prediction framework posits that people continuously predict potential changes in their shared environments, including social information about each other, and the interdependence of interacting people’s predictions result in increased INS (Bilek et al., 2022; Kingsbury et al., 2019; Mayo & Shamay-Tsoory, 2024). A further strength of the mutual prediction framework is that it offers a theoretical basis for differentiating between within-brain connectivity (i.e., functional connectivity) and between-brain connectivity (i.e., INS). The distinction is particularly informative in asymmetric interactions (Heggli et al., 2019, 2021; Mayo & Shamay-Tsoory, 2024). For example, in leader–follower settings, a computational model shows greater within-brain connectivity in the leader than the follower (Heggli et al., 2019), reflecting how the follower adapts to the leader unilaterally. By contrast, when two individuals adapt to each other mutually without defined turn-taking patterns, both show greater between-brain connectivity than within-brain connectivity (Heggli et al., 2019, 2021). Adding nuance to this explanation, Mayo and Shamay-Tsoory (2024) present a computational model of leader–follower interaction with a focus on the internal models that would support each role. The findings describe leaders predicting their own behaviour and followers predicting their partner’s behaviour. Evidence further suggests that within-brain and between-brain connectivity are associated with different functional demands in the emergence of behavioural synchrony: Within-brain connectivity plays a role in the early phases of initiating behavioural synchrony, whereas between-brain connectivity supports the maintenance of synchronised behaviour once it is established (Marton-Alper et al., 2023).

Considered together, these findings highlight how the neural dynamics of interacting partners, just as the dynamics within a single brain, are complex and oscillate between patterns of segregation and integration that are shaped by the functional demands of the task (Heggli et al., 2019; Marton-Alper et al., 2023; Park & Friston, 2013; Shamay-Tsoory, 2022). In other words, the different dynamics (e.g., individual action, mutual or asymmetric adaptation, and turn-taking or simultaneous coordinated behaviour) shape both the strength and configuration of functional connectivity within and between brains. A closer examination of these configurations, embedded in the mutual prediction framework and specifically taking spatial and temporal information into account, can inform our understanding of social interactions (Mayo & Shamay-Tsoory, 2024).

### 1.3 Two-brain states as a tool for exploring interpersonal dynamics

When computing INS levels from two people’s fNIRS signals (the technique employed in this study), researchers commonly use a signal processing method called wavelet transform coherence (WTC; see Hakim et al., 2023 for detailed review of INS metrics). WTC returns the local correlation between two signals for a given channel pair in a given time and frequency window (Gvirts et al., 2023). The extracted values are then typically averaged across the task duration. When applied to signals from two interacting individuals, WTC can be interpreted as an extension of conventional functional connectivity, operationalizing correlation-based dependency between neural timeseries, but across brains rather than within one brain (Hasson et al., 2012).

Averaging WTC values across an interaction between two individuals per region might however obscure the temporal dynamics of INS. Insights into the temporal dynamics of INS are especially relevant for tracking and interpreting INS in naturalistic settings where people build relationships across multiple sessions and where the interaction dynamics are not externally paced by a pre-specified experimental design. Temporal dynamics of INS can be preserved using approaches that quantify changes in INS at a biologically relevant timestep (e.g., seconds for haemodynamic fluctuations recorded with fNIRS).

The spatial dynamics of INS are also typically obscured via commonly used unidimensional metrics of INS (e.g., computing WTC between channel-pair, then conducting comparisons at the group level per channel pairing, which are then visualised to report as findings). This can be overcome by incorporating the spatial mapping of the INS values (e.g., as a matrix rather than a single value) into the analysis before conducting comparisons at the group level. Introducing the spatial dimension to dyad-level analysis can shed light on the configurations of functional connectivity within and between brains.

One way of conceptualising INS analyses that preserve temporal and spatial dynamics is to identify recurrent configurations of functional connectivity, i.e., *brain states*, in the brain signals recorded from interacting dyads. This can be achieved by extending the identification of brain states from within single brains to within and between dyads’ brains, wherein both brains are considered components of a single integrated system (Shamay-Tsoory, 2022). Li and colleagues (2025) offer a recent proof of concept that this approach can be conducted in EEG data, by extending the concept of microstates, quasi-stable configurations of brain activity (Michel & Koenig, 2018), to joint analysis of dyadic neural data. Their analysis revealed that two-brain microstates differed between observer—actor paradigms and tasks with more symmetrical demands (Q. Li et al., 2025). With this approach, Li and colleagues were able to examine concurrent patterns of brain activity but did not measure functional connectivity. Two-brain states of connectivity can be used to investigate how regions within and across brains are integrated or segregated over time.

To identify brain states in fNIRS data, some studies used *a priori* definitions (Lu et al., 2022) or behavioural markers (Dai et al., 2024), though these constrain the space of possible states to those anticipated in advance. Focusing on fluctuations in functional connectivity (i.e., dynamic functional connectivity) in fNIRS data, Zhang & Zhu (2020) used a *k*-Means clustering analysis on Pearson correlation matrices of single-brain activity in time windows. They show that such an approach can be useful to reduce the complexity of high-dimensional connectivity data while respecting the spatial configuration and temporal dynamics by replicating previous results using other brain imaging techniques (Y. Zhang & Zhu, 2020). Li and colleagues (2021) present a two-brain extension of this approach, clustering channel-pair WTC matrices derived from time windows in fNIRS data collected during a dyadic creative design task. This provides novel insights into the neural dynamics of social interactions; however, their analysis focuses on inter-brain channel pairs. The full bandwidth of two-brain states may include patterns that are characterised as much by the extent of within-brain connectivity as by the extent of between-brain connectivity (Heggli et al., 2019; Marton-Alper et al., 2023; Mayo & Shamay-Tsoory, 2024).

Integrating information on both is therefore necessary to gain the full picture. Näher and colleagues (2024) further demonstrated that the usage of additional measures of similarity between neural signals in tandem with Riemannian geometry-based approaches that preserve the intrinsic structure of inter-channel relationships outperforms traditional models in classification of brain states. These findings motivate an approach that applies Riemannian geometry-based clustering models to within-brain and between-brain dynamic functional connectivity data, enabling the discovery of two-brain states in a data-driven manner.

### 1.4 Present study: Exploring intergenerational collaborative interactions using two-brain states

This preregistered study (<u>anonymised link</u>) is a secondary analysis of an existing fNIRS hyperscanning dataset (<u>anonymised link</u>). We applied two-brain state analysis approach to the dataset, which includes 732 individual fNIRS recordings from 61 dyads who each attended 6 creative drawing sessions. Thirty of the dyads are same generation dyads (young adults) and 31 are intergenerational (1 young adult with 1 older adult).

To gain insight into the symmetric and asymmetric interaction dynamics involved in intergenerational collaborative drawing, we aimed to:

1. Identify two-brain states based on dynamic functional connectivity in fNIRS data acquired during collaborative drawing in IFG and TPJ.
2. Examine the extent to which collaborative vs. independent activity and same generation vs. intergenerational interactions give rise to specific two-brain states.
3. Investigate the extent to which any identified two-brain states change across repeated encounters.

Analyses of combined single-brain states were also included in our preregistration and are presented in the Supplementary Material. In non-preregistered analyses, we also explored the extent to which turn-taking behaviour occurs during same generation and intergenerational collaborative drawing, and the extent to which turn-taking and specific two-brain states are associated.

## 2 Methods

### 2.1 Transparency and openness

This study consists of secondary analysis of the fNIRS data, from which initial analyses of INS levels during same generation and intergenerational collaborative drawing are presented by [redacted for peer review]. The entirety of this longitudinal hyperscanning dataset will be shared publicly in the near future [redacted for peer review]. Our pre-registered data analysis plan as well as data and code can be found on the Open Science Framework (OSF; <u>anonymised link</u>). We report how we determined our sample size, all data exclusions (if any), all manipulations, and all measures in the study (Munafò et al., 2017).

### 2.2 Participants

We aimed to collect as many useable datasets as possible between November 2023 and June 2024. The final sample consists of 122 participants. We assigned older adults (*n* = 31, age 70+ years) and younger adults (*n* = 91, age 18–35 years) to same generation or intergenerational dyads. Same generation dyads (*n* = 30) consisted of only younger adults (22.5 years [SD = 3.6]; 36 female, 24 male), intergenerational dyads (*n* = 31) consisted of one older adult (76 years [SD = 4];18 female, 13 male) and one younger adult (24 years [SD = 4]; 19 female, 11 male, 1 other). Same generation dyads had a mean age difference of 3.5 years (SD = 3.4) and intergenerational dyads had a mean age difference of 52.1 years (SD = 5.9). All dyads began the study as strangers; they had never met prior to the first experimental session.

We recruited participants through [redacted for peer review] university recruitment pool, the [redacted for peer review] University center, as well as local seniors’ clubs and community organizations (e.g., choirs, theatre groups, and orchestras), via social media and paper flyers. We included individuals who spoke German fluently and did not report having any known psychological or neurological disorders (e.g., stroke, ADHD, depression) when completing the screening questionnaire.

We obtained written informed consent from all participants. Ethical approval was obtained from the Ethics Committee of the [redacted for peer review] (Ref: 2023-01073). The study was conducted according to the Declaration of Helsinki.

### 2.3 Procedure

#### Experimental sessions

Each dyad attended six experimental sessions, spaced roughly one week apart. Upon arrival, participants completed questionnaires, separated from one another by a felt partition wall for privacy. The questionnaires measured loneliness levels (De Jong Gierveld & van Tilburg, 2006) and attitudes toward individuals from their own and the other (older or younger) generation (Pittinsky et al., 2011). Participants completed an empathy questionnaire at the first session only (Paulus, 2009). Next, fNIRS caps were placed on participants’ heads and images for 3D head models were acquired. Then, fNIRS sensors were positioned on caps and fNIRS devices turned on, signal quality was optimised by additional hair clearing, and fNIRS recordings were initiated.

At each session, dyads completed three 5-min blocks of drawing with oil pastels. First, dyads drew alone for 5 min on separate pieces of paper, separated from one another by a felt partition wall so they could not see each other. Next, the partition wall was removed. Dyads received a piece of A3 art paper, on which to draw together for 5 min, twice. To ensure constant levels of conversation between drawing conditions (i.e., alone and together), we instructed participants that they should not talk while drawing, and that they could talk as much as they desired during the rest of the experimental session. We provided no instructions as to what or how to draw, allowing participants and dyads maximal freedom to interact freely in a naturalistic environment.

Following the final drawing block, dyads then completed a short collaborative task that varied each week to maintain motivation across sessions (e.g., speed-based tasks, a jigsaw puzzle, a divergent thinking task). The felt wall was placed between participants, who then completed a measure of social closeness (Aron et al., 1992) and wrote subjective comments on the experimental session. After the final session, the lead experimenter conducted a semi-structured interview with each participant about their experience in the study.

The data analysed in this study consists of the fNIRS recordings during the three drawing blocks.

#### fNIRS acquisition

To record cortical hemodynamic activity during the drawing and collaborative tasks, we used two Cortivision Photon Cap fNIRS systems with a sampling frequency of ~5 Hz. We positioned 26 optodes (16 sources emitting wavelengths of 760 and 850 nm, and 10 detectors) over four regions of interest (ROIs): the left and right inferior frontal gyrus (IFG) and the left and right temporoparietal junction (TPJ). The optode montage is visualised in Supplementary Figure 1. To synchronise the fNIRS recordings, we used PsychoPy (Peirce et al., 2022) to write triggers marking the beginning of each drawing and the collaborative activity directly into the recordings.

#### Optode digitisation

After the researcher placed the fNIRS cap on a participant’s head and before securing the optodes, the researcher recorded a 3D head scan using Structure Sensor Pro connected to an Apple iPad Pro.

Using MATLAB and the Fieldtrip toolbox (Oostenveld et al., 2011), we manually co-registered the head scan to MNI space. We then used optode positions in MNI coordinates to calculate channel locations in MNI space, inter-optode distances, and distances between channel locations and centres of regions of interest. We calculated these distance measures to assess channel quality and positional stability across sessions.

#### Quantification of turn-taking

We assume that mutual prediction is stronger during simultaneous drawing, where both participants draw and observe the other person drawing, than during drawing with overt turn-taking, where one participant observes and the other draws. To quantify turn-taking (or absence thereof) while dyads drew together, the lead experimenter scored the extent to which dyads took turns drawing and drew simultaneously. The experimenter watched a timer counting down from 5 minutes and estimated the percentage of the 5 minutes that dyads drew simultaneously in increments of 10%. The percentage was then converted into a score between 0 (reflecting 100% turn-taking and 0% simultaneous drawing) and 10 (reflecting 0% turn-taking and 100% simultaneous drawing). To invert the scale, we subtracted scores from 10, so 0 can be interpreted as no turn-taking (only simultaneous drawing), and 10 can be interpreted as purely overt turn-taking (no simultaneous drawing).

### 2.4 Data analysis

#### Missing data

Though we endeavoured to include fNIRS recordings characterizing all combinations of participants, sessions, and ROIs, some data were missing or excluded from the analysis due to technical issues (e.g., signal interruptions caused by Bluetooth failure) or low data quality identified during the channel selection process (described below). Data exclusions are detailed in Supplementary Table 1.

#### fNIRS channel selection

We retained one channel per ROI to the following criteria, to ensure the inclusion of high-quality signals, as previously done by De Felice and colleagues (2024). Included channels showed visual evidence of cardiac oscillation, inter-optode distance between 17 mm and 40 mm (Brigadoi & Cooper, 2015) and scalp coupling indices (SCI) > 0.7 and <1. The SCI was calculated in *MNE-nirs* (Luke et al., 2021) by cross-correlation between the signal at the two wavelengths between 0.7 and 1.35 Hz where cardiac oscillation is expected (Pollonini et al., 2014). To select the best channel for a given participant, session and ROI, we used the channel physically located closest to the centre of the region of interest when more than one channel survived the inclusion criteria above.

#### fNIRS preprocessing

Using *MNE-python* (Gramfort et al., 2014) and *MNE-nirs* (Luke et al., 2021), we first resampled the raw optical intensity signals to 5 Hz. We converted signals to optical density and corrected Motion artifacts using Temporal Derivative Distribution Repair (Fishburn et al., 2019) to remove extracerebral systemic noise like cardiac, respiratory, baroreceptive, or sympathetic activity (Yücel et al., 2021), we conducted short channel regression using the short channel with the highest SCI. We then converted signals to oxygenated and deoxygenated haemoglobin concentration values using the modified Beer–Lambert law and an age-adjusted partial pathlength factor (Garcia-Castro, n.d.; Scholkmann & Wolf, 2013). This age-adjustment is validated for ages 0-70 years. We set the age parameter to 70 for participants >70 years. We filtered the signals using a band-pass filter between 0.015 and 4 Hz. Next, we segmented signals for each channel into task blocks (i.e., drawing alone, drawing together) and z-standardised segmented signals within channels.

#### Pseudo dyads

To validate whether the properties of identified brain states are specific to true interaction, we compared real dyads with pseudo dyads, generated from all combinations of participants who did not interact as a pair in the dataset. Pseudo dyads included same generation dyads of two younger adults and intergenerational dyads, but no same generation dyads of two older adults.

### 2.5 Identifying brain states and extracting brain-state timeseries

We performed the following computations on the [redacted for peer review] high-performance computing cluster at [redacted for peer review], using *scikit-learn* (Pedregosa et al., 2011), and *pyriemann* (Barachant et al., 2026). The clustering pipeline consisted of five key components (Figure 1). First, we concatenated individuals’ signals into dyadic (two-brain) data frames to represent dyads (real and pseudo) as a unit. Second, we temporally segmented the dyadic signals using a sliding window. Third, we estimated neural connectivity matrices mapping within-brain and between-brain connectivity using four kernel functions (i.e., similarity metrics) on the temporally segmented signals. Fourth, we identified brain states using a clustering algorithm. Fifth and finally, we assigned each 15-s signal segment to its respective brain state, generating a time series of brain states.

**Figure 1.**
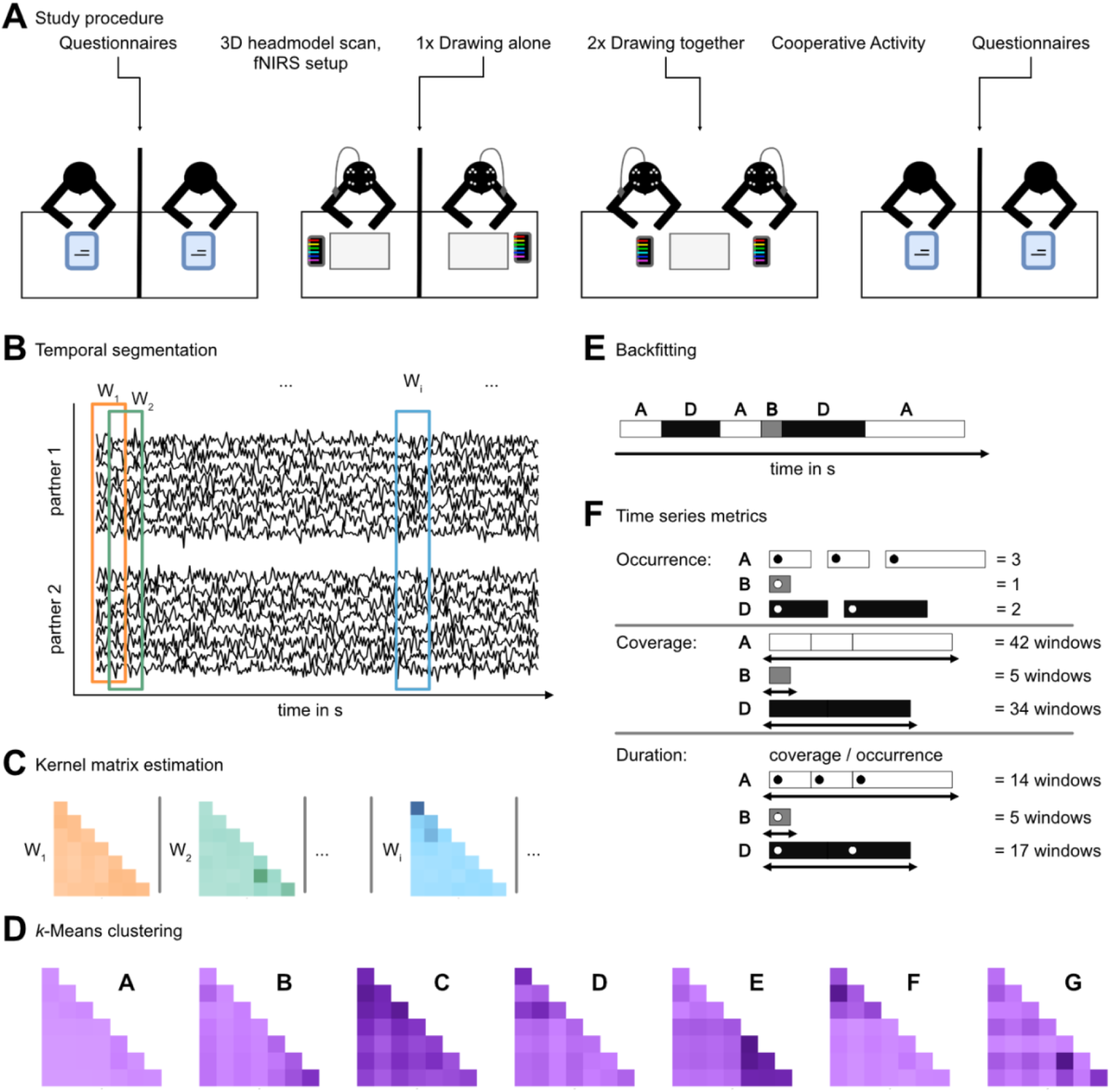
A) From left to right, a summary of the protocol followed at each session. B) The time series of fNIRS signals recorded from 2 dyad members were concatenated into a single dataframe. Coloured boxes represent discrete windows (W_*i*_) for which kernel matrices were computed. Window length was 15 s and windows were shifted in increments of 1 s. C) Using 5 different kernel functions, kernel matrices were computed and shrunk per window for HbO and HbR separately. Per window, kernel function, and shrinkage parameter, the HbO and HbR matrices were combined into a block diagonal matrix. D) Using block diagonal matrices of real dyads only, a Riemannian-geometry-based k-Means identified recurrent brain states. Optimal hyperparameters (kernel function, shrinkage, number of clusters) were determined in a brute-force grid search. E) The identified states were then backfit onto time series of real and pseudo dyads, such that each window received a state label. F) The temporal dynamics of the two-brain states per real and pseudo dyad and per task were quantified using 3 metrics, namely occurrence, duration, and coverage. Group level statistical comparisons were conducted on these metrics.

#### Concatenation

Per dyad and task block, we concatenated the fNIRS signals (8-channel timeseries) concatenated to create a 16-channel timeseries. Within dyads, concatenation order was fixed, such that the same dyad members’ signals are first in each concatenated time series. This is specifically relevant for intergenerational dyads, where the younger dyad member’s signals (channels 1-8 of joint timeseries) preceded the older dyad member’s signals (channels 9-16 of joint timeseries).

#### Temporal segmentation and kernel matrix estimation

Per dyad and task block, timeseries were segmented using a sliding window following Li and colleagues (2021) with window size set to 15-s and step size to 1 s. We selected five kernel functions to quantify similarity between neural signals in a time window, which is returned as a kernel matrix. Specifically, we selected a subset of the kernel function used by Näher and colleagues (2024), prioritising kernel functions with unique properties. The selected kernel functions included covariance, correlation coefficient, Tyler’s M-estimator-based covariance, Ledoit–Wolf shrunk covariance, and the radial basis function kernel (further detailed in the Supplementary Material). To ensure positive definiteness and numerical stability for clustering algorithms, we applied additional kernel-level shrinkage to all resulting kernel matrices (Upton & Cook, 2014). As part of hyperparameter tuning, the shrinkage parameter λ was set to three different values; details of the tuning procedure are provided below. For each shrinkage parameter, kernel function, and time window, we derived and shrunk kernel matrices separately for oxygenated haemoglobin (HbO) and deoxygenated haemoglobin (HbR), and then combined them into a block diagonal matrix (Näher et al., 2024). For readers interested in gaining familiarity with kernel matrix estimation, we recommend perusing Shawe-Taylor and Cristianini (2004). Each 15-s window is transformed into kernel matrices, then the window is shifted by 1 s and the new resulting window is transformed into new kernel matrices.

#### Clustering procedure

Based on a recent demonstration that Riemannian geometry-based models outperform classical models in classification of fNIRS brain states (Näher et al., 2024), we applied a Riemannian geometry-based *k*-means clustering algorithm to identify brain states during activities. Clusters were identified using data from real dyads only. To determine which kernel function and shrinkage parameter to employ (i.e., via hyperparameter tuning), we employed a brute-force grid search. We entered the combined block diagonal matrices containing HbO and HbR connectivity into clustering and subsequent hyperparameter tuning. The hyperparameter space is detailed in Supplementary Table 3. We determined the best combination of hyperparameters using the Calinski-Harabasz index (CHI; Calinski & Harabasz, 1974), which favours coarser *k*-Means solutions with strong global separation, and the Davies-Bouldin index (DBI; Davies & Bouldin, 1979), which favours finer *k*-Means solutions that are locally well defined. The CHI indicated a three-cluster solution, while the DBI indicated a seven-cluster solution. Further selected hyperparameters are reported in Supplementary Table 2. Here, we report the seven-cluster solution favoured by DBI which was more detailed and therefore more informative for subsequent analyses and interpretation; the three-cluster solution is reported in the Supplementary Material.

#### Metrics of temporal dynamics of two-brain states extracted from brain-state timeseries

After hyperparameter tuning, we fit back the clustering with the best combination of hyperparameters into the data sets. Each 15-s window is assigned a state, yielding a time series of brain-state sequences. We estimated three metrics commonly used to describe state sequence time series: occurrence, coverage, and duration. *Occurrence* refers to the number of discrete entries into a state during the task (consecutive windows in a given state are counted as one instance). *Coverage* refers to the total number of windows in which a state is active. *Duration* is the mean number of windows in which a given state is active consecutively and can thus be interpreted as a measure of a given state’s stability.

### 2.6 Statistical analyses

The primary aim of this study was to explore the extent to which dyadic brain states differ between collaborative and independent drawing (drawing together vs. drawing alone). In addition, we explored the influence of dyad composition (intergenerational vs. same generation, hereafter called group). In addition to our pre-registered exploratory analyses of coverage, we also conducted non-preregistered exploratory analyses of occurrence and duration, and the relationship between turn-taking and a specific state identified as differing between groups. We carried out the analyses below using R and the R packages *emmeans* (Lenth, 2021), *lme4* (Bates et al., 2015), *lmtest* (Zeileis & Hothorn, 2002), *MASS* (Venables & Ripley, 2002), *purrr* (Wickham & Henry, 2026), and *tidyverse* (Wickham et al., 2019). We made graphics using the packages *ggplot2* (Wickham, 2016) and *viridis* (Garnier et al., 2023).

To establish whether each brain state was the result of true dyadic interactions or could also be observed when the signals of who did not interact were aligned, we contrasted real and pseudo dyads. We fit separate negative binomial models for each metric (n = 3; rationale described in Supplementary Material). The formula was the following: *metric ~ state * drawingTask * dyadType * group * session*, where drawingTask refers to drawing together or alone, dyadType refers to real and pseudo dyads, and group refers to same generation and intergeneration groups. We identified brain states showing significant differences between real and pseudo dyads using planned contrasts for each combination of task and group. In the case that any single combination (e.g., drawing alone and same generation group) differed significantly between real and pseudo dyads for a given brain state, we conducted further contrasts between task, group, and the interaction thereof, on that brain state. To account for multiple comparisons, *p*-values in contrast analyses were adjusted using the False Discovery Rate (FDR) procedure (Benjamini & Hochberg, 1995). The same process was repeated for the slope of each metric across sessions.

Finally, we explored the relationship between the duration of a state that differs between groups, while dyads draw together, and turn-taking behaviour. We fit a general linear model with the following formula: *turn taking ~ duration * group + session*. Influential observations were excluded prior to final model estimation if their Cook’s distances exceeded the conventional threshold of *4/n* (Bollen & Jackman, 1990). Though this exploratory analysis was not preregistered for this study, the collection of the turn-taking measure was included in the pre-registration of the primary dataset (<u>link anonymized for peer review</u>).

## 3 Results

First, we identified the two-brain states during creative drawing, based on the combined HbO and HbR block diagonal matrices. We set qualitative thresholds based on the distributions of connectivity values to describe the strength of connections: high >= 0.3, medium >= 0.25, low >= 0.2, very low < 0.2. We identified seven two-brain states, which are labelled with letters from A to G, are visualised in Figure 2. States A, B, and C correspond to relatively low, medium, and high connectivity within and between brains, respectively (i.e., A = low, B = low to medium, C = high). States D and E correspond to low-to-medium connectivity between brains and medium-to-high connectivity within only one dyad member’s brain (D for first and E for second dyad member). States F and G correspond to low-to-medium connectivity between brains and prominent within-brain connectivity specifically between bilateral IFG in only one dyad member (F for first and G for second dyad member).

**Figure 2.**
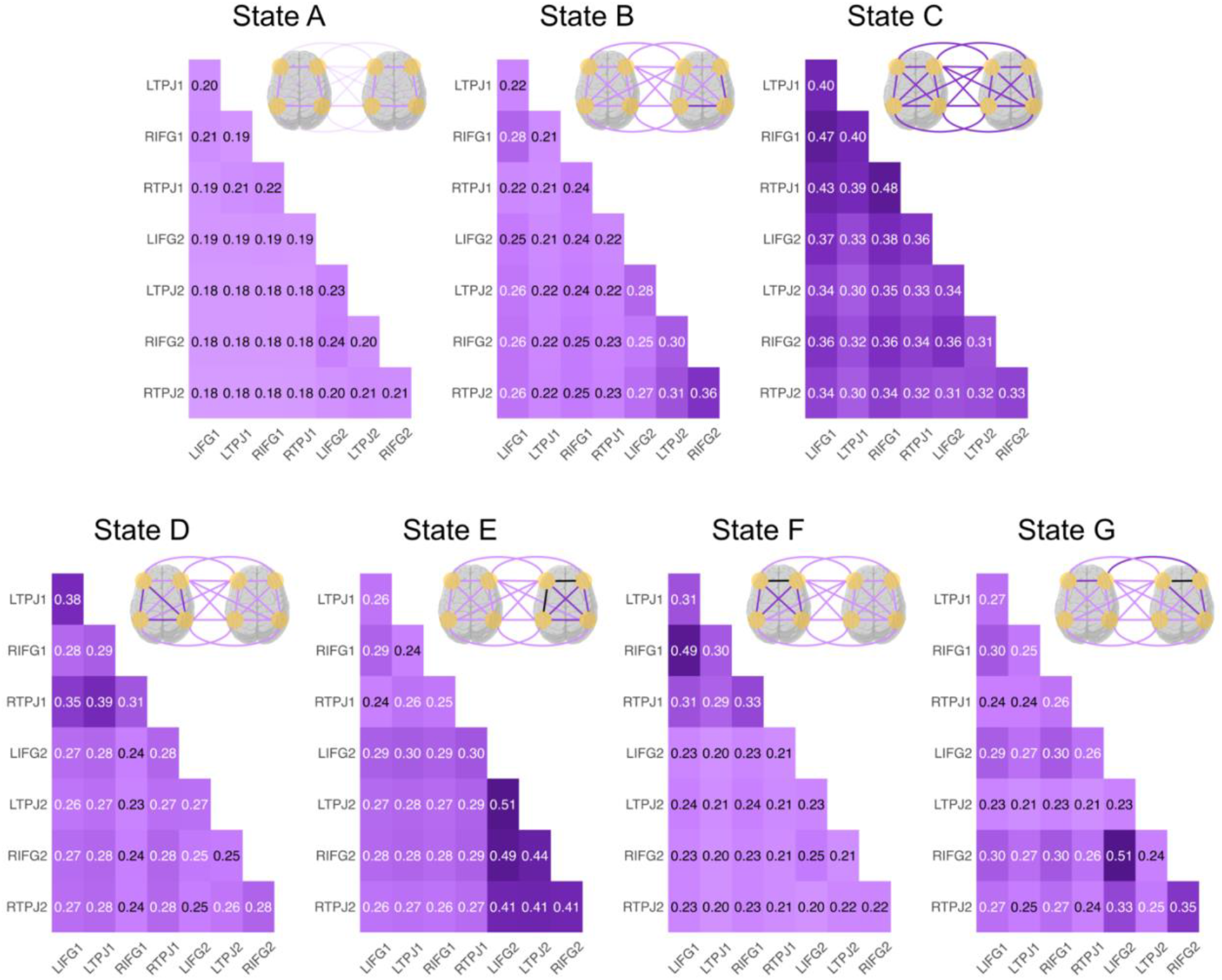
Seven two-brain states identified using Riemannian geometry-based clustering of dynamic functional connectivity matrices. Matrices show similarities in the HbO signals between ROIs belonging to each dyad member (distinguished by 1 and 2). States A-B show low, medium, and high connectivity within and between brains, respectively. States D-E show low-to-medium connectivity between brains and high connectivity within only one dyad member’s brain. States F-G show low-to-medium connectivity between brains and prominent within-brain connectivity, specifically between bilateral IFG in only one dyad member. The pairs of brains with purple lines show the strength of the within and between-brain connectivity; lighter purple = lower connectivity, darker purple = greater connectivity. Connectivity matrices for HbR signal show high levels of similarity to these matrices and are visualised in Supplementary Figure 2.

For each brain-state and metric (occurrence, coverage, duration) and brain-state (A–G) we first assessed the extent to which the metric differs between real and pseudo dyads. Next, using planned contrasts, we evaluated the extent to which dyadic brain states differ between collaborative and independent drawing, intergenerational and same generation dyads, and the interaction between these contexts.

### 3.1 Occurrence (number of discrete entries into a brain state)

#### Real vs. pseudo dyads

We observed differences between real and pseudo dyads for state B (Intergenerational group, drawing together: *z* = -3.29, *p*_adj_ = .014) and state D (Intergenerational group, drawing alone: *z* = 3.39, *p*_adj_ = .014). States A, C, E, F and G did not differ significantly between real and pseudo dyads for any drawing condition or group (Supplementary Tables 4 and 5).

#### States B and D

Likelihood ratio tests showed main effects of state and task (*p* < .05; Supplementary Table 6). Contrast analyses revealed no statistically significant differences in any states between tasks or groups after FDR correction. There were a trend towards lower occurrence of state D in drawing alone compared to drawing together, but only in intergenerational dyads (*z* = -2.62, *p*_adj_ = .053), and a trend towards lower occurrence of state D in intergenerational dyads compared to same generation dyads, but only for drawing alone (*z* = -2.72, *p*_adj_ = .053; Figure 3; Supplementary Table 7).

**Figure 3.**
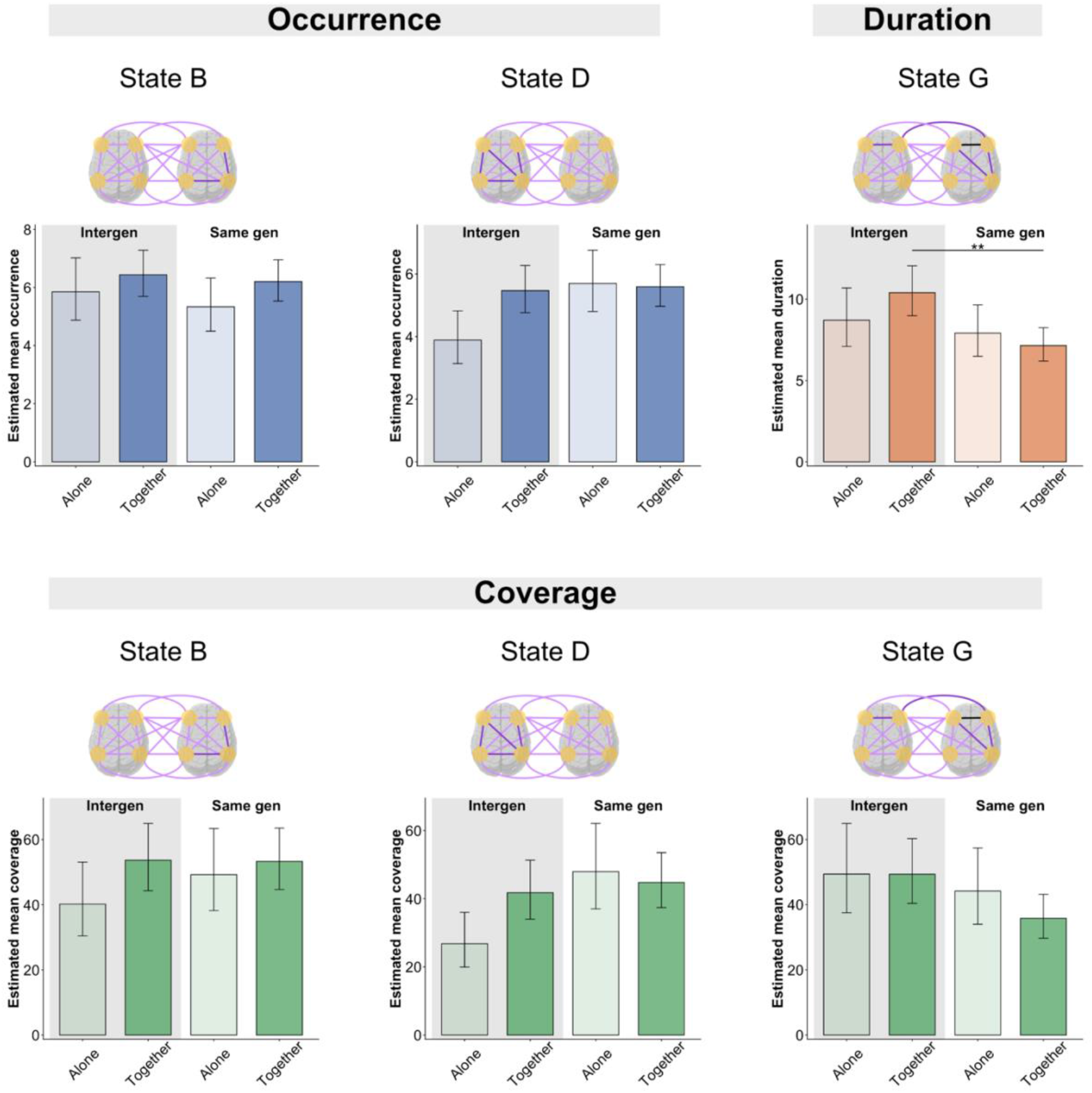
Per metric, the two-brain states that differed significantly between real and pseudo dyads. Planned contrasts showed no significant difference between task, group, or task*group interactions for occurrence or coverage. For duration, a significant difference between groups was observed for drawing together. As in Figure 2, the pairs of brains with purple lines show the strength of the within and between-brain connectivity for each state; lighter purple = lower connectivity, darker purple = greater connectivity.

### 3.2 Coverage (total number of windows in which a brain state is active)

#### Real vs. pseudo dyads

We observed differences between real and pseudo dyads for state B (Intergenerational group, drawing together: *z* = -2.98, *p*_adj_ = .027), state D (Intergenerational group, drawing alone: *z* = 3.18, *p*_adj_ = .020), and state G (Intergenerational group, drawing together: *z* = -3.30, *p*_adj_ = .020). We observed a trending difference for state F (Intergenerational group, drawing together: *z* = 2.68, *p*_adj_ = .052). States A, C, and E did not differ significantly between real and pseudo dyads for any drawing condition or group (Supplementary Tables 8 and 9). No further analyses were conducted for state F.

#### States B, D, G

Likelihood ratio tests showed a main effect of state, as well as interaction effects for state and group, and task and group (*p* < .05; Supplementary Table 10). Contrast analyses revealed no statistically significant differences in any states between tasks or groups after FDR correction. There were trends towards higher coverage of state D during drawing together than alone, but only for intergenerational dyads (*z* = 2.41, *p*_adj_ = .095), as well as towards lower coverage of state D (*z* = -2.90, *p*_adj_ = .067) and higher coverage of state G (*z* = -2.30, *p*_adj_ = .097) in the intergenerational group compared to the same generation group, but only while drawing alone (Figure 3; Supplementary Table 11).

### 3.3 Duration (mean number of consecutive windows in which a brain state is active)

#### Real vs. pseudo dyads

We observed a difference between real and pseudo dyads for state G (Intergenerational group, drawing together: *z* = –4.51, *p*_adj_ < .001). States A–F did not differ significantly between real and pseudo dyads for any drawing condition or group (Supplementary Tables 12 and 13).

#### State G

Likelihood ratio tests showed a significant effect of group (*p* < .05; Supplementary Table 14). Contrast analyses revealed a significant difference while drawing together, where intergenerational dyads showed significantly longer durations than same generation dyads (*z* = 3.59, *p*_adj_ = .002; Figure 3; Supplementary Table 15).

### 3.4 Longitudinal changes in two-brain states

We examined the differences in the slopes of occurrence, coverage, and duration of the two-brain states across sessions between real dyads and pseudo dyads. There were no significant differences in occurrence, duration or coverage after FDR correction (Supplementary Tables 16–18), therefore, we did not conduct any further analyses.

### 3.5 Turn-taking during collaborative drawing

During the review process, insightful questions raised by reviewers led us to explore the relationship between the duration of state G and turn-taking behaviour. We fit an exploratory non-preregistered general linear model, which indicated a group difference in the amount of turn-taking behaviour while dyads drew together: intergenerational dyads engaged in more turn-taking behaviour than same generation dyads (*b* = -1.602, *SE* = 0.492, *t* = -3.254, *p* = .001; Figure 4A). In other words, same generation dyads engaged in simultaneous drawing more often than intergenerational dyads. Turn-taking behaviour remained stable across sessions (*b* = -0.111, *SE* = 0.093, *t* = -1.201, *p* = .232; Figure 4A). We also found that the duration of state G was significantly associated with turn-taking behaviour, such that longer durations were associated with less turn-taking (*b* = -0.069, *SE* = 0.023, *t* = -3.051, *p* = .003; Figure 4B). We did not observe a significant difference in the slope of this relationship between intergenerational and same generation dyads (Δ*b* = -0.028, *SE* = 0.056, *t* = -0.498, *p* = .619).

**Figure 4.**
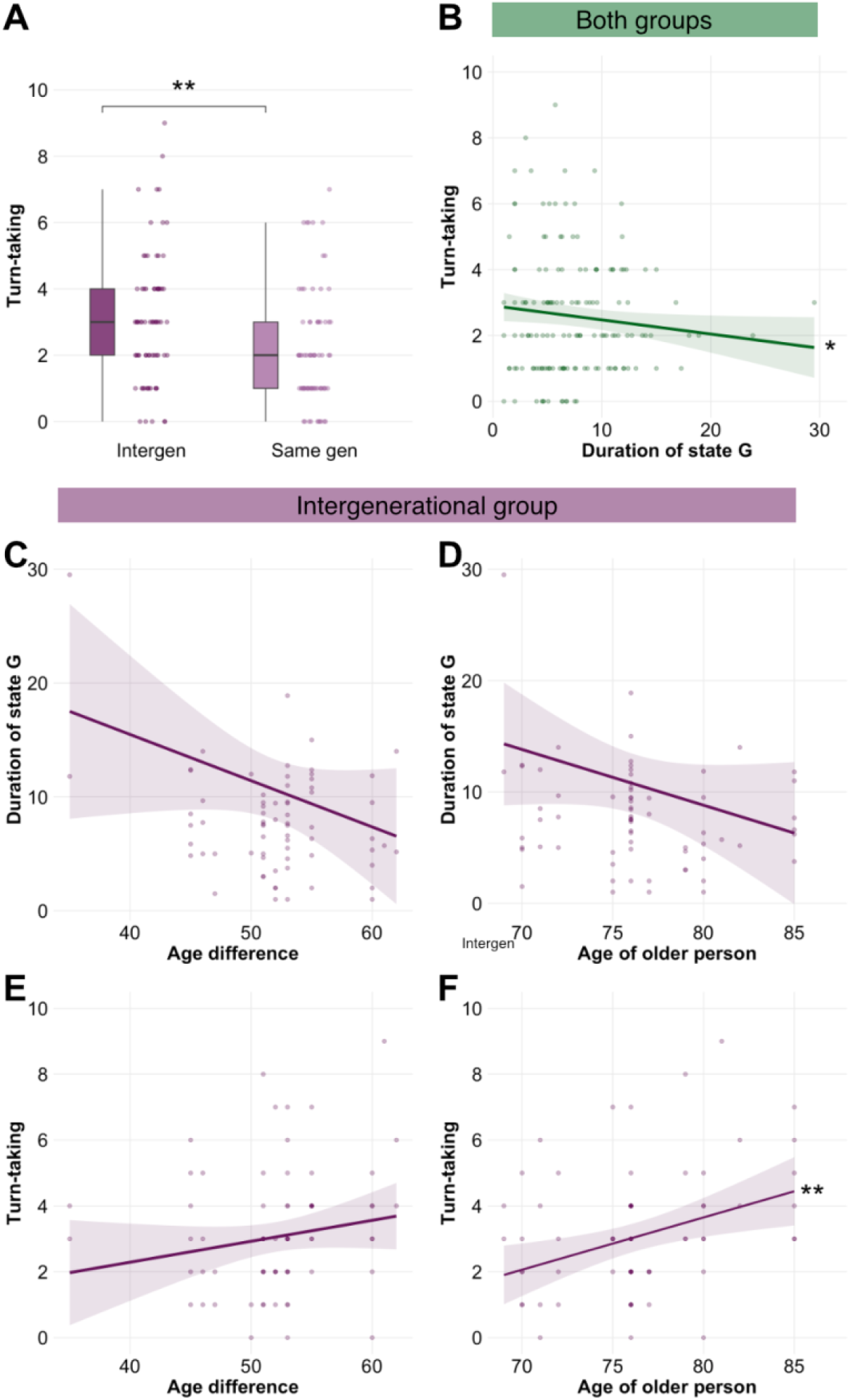
A) Turn-taking behaviour while drawing together. The researchers estimated the percentage of the 5 minutes that dyads drew simultaneously in increments of 10%. Zero can be interpreted as no turn-taking (only simultaneous drawing), and 10 can be interpreted as purely overt turn-taking (no simultaneous drawing). B) The significant negative relationship between duration of state G and turn-taking behaviour while dyads drew together (groups aggregated). C) The non-significant (numerically negative) relationship between duration of state G and age difference in intergenerational dyads. D) The non-significant (numerically negative) relationship between duration of state G and the age of the older dyad member in intergenerational dyads. E) The non-significant (numerically positve) relationship between turn-taking and age difference in intergenerational dyads. F) The significant positive relationship between turn-taking and the older dyad member’s age in intergenerational dyads.

### 3.6 Age differences influence state G duration and turn-taking

During the review process, we also reflected on the role of older age in shaping the duration of state G and turn-taking behaviour. Our planned analyses established that the duration of state G differs significantly between real and pseudo intergenerational dyads (Section 3.3), validating that state G is related to real interactions rather than the presence of an older adult in a dyad (elaborated upon in Discussion).

Focusing on intergenerational dyads drawing together, we fit non-preregistered and exploratory models to examine the extent to which age differences and the age of the older dyad member may influence the duration of state G in intergenerational dyads, and in separate models, turn-taking behaviour (Figure 4C-F; Supplementary Table 37 to 40). We acknowledge that age differences and the age of the older dyad member are inherently non-independent and provide results from both in the spirit of hypothesis generation. For the duration of state G, the models returned numeric trends, but no statistically significant relationships, where larger age differences (*b* = -0.406, *SE* = 0.267, *t* = -1.519, *p* = .133; Figure 4C) and greater age of the older dyad member (*b* = -0.501, *SE* = 0.333, *t* = -1.502, *p* = .138; Figure 4D) co-occur with shorter durations of state G. For turn-taking behaviour, the models show that age differences do not significantly predict turn-taking (*b* = 0.064, *SE* = 0.045, *t* = 1.405, *p* = .165; Figure 4E), while the older dyad member’s age is significantly and positively associated with turn-taking (*b* = 0.159, *SE* = 0.054, *t* = 2.940, *p* = .004; Figure 4F).

## 4 Discussion

In this study, we implemented an approach to characterise shared patterns of neural activity during a type of naturalistic social interaction that captures richer spatial and temporal information than traditional approaches. Specifically, we identified seven distinct two-brain states, three of which were unique to real dyads, as opposed to pseudo dyads. Of these three two-brain states, only one (G) differed significantly in duration between same and intergenerational dyads, and only while dyads drew together. The increased duration of state G, which involved medium-to-low between-brain connectivity and within-brain bilateral IFG connectivity for one dyad member, likely reflects an intergenerational collaboration pattern, where the older person leads and the younger person follows. Longer durations of state G were associated with less turn-taking behaviour (i.e., more simultaneous drawing). No significant longitudinal changes in the temporal dynamics of any two-brain states were observed across the six drawing sessions. We explore the implications of these findings below.

### 4.1 Symmetrical within- and between-brain connectivity may reflect common cognitive processing

The first group of states A-C shows symmetrical patterns of connectivity (Figure 2). These states show low, medium, and high connectivity within and between brains, respectively. From a theoretical perspective, the observed patterns do not align with the mutual prediction framework, where continuous prediction of a partner’s behaviour during the collaborative drawing should result in divergent patterns of intra- and interbrain synchrony (Mayo & Shamay-Tsoory, 2024). Instead, this pattern could reflect common cognitive processing, i.e., that both dyad members’ brains are responding similarly to the same stimulus (Burgess, 2013). Taking this perspective, the different strengths (low, medium, and high) of connectivity would reflect the degree to which the dyad members perceive the same stimuli differently. However, only state B differed between real and pseudo dyads, specifically occurring less often for real than pseudo dyads when intergenerational dyads drew together, and did not differ significantly from other group/task combinations. In other words, none of the identified symmetrical two-brain states of varying strengths were specific to a group or task in real dyads. This is perhaps not surprising, as the open-ended nature of the creative task, drawing alone or together in silence, is unlikely to result in both dyad members perceiving the same stimuli the same way at the same time, as might happen if both individuals view the same visual stimuli or complete a clearly structured task (Finn et al., 2020; R. Li et al., 2021). Moving away from common cognitive processing, an alternative explanation for fleeting symmetrical two-brain states may be that they reflect transitions between less symmetrical two-brain states (Hendrikse et al., 2025; Mayo & Gordon, 2020; Roche et al., 2025).

### 4.2 Asymmetric connectivity patterns during independent activity unlikely to reflect leader–follower dynamics

States D and E feature low-to-medium connectivity overall, with prominent within-brain connectivity in one dyad member (Figure 2). State D, but not E, differed in occurrence and coverage between real and pseudo dyads. This emerged specifically for same generation dyads when drawing alone. As same generation dyad members are virtually indistinguishable, the relevance of the arbitrarily assigned first dyad member showing greater within-brain connectivity is not immediately clear. Our planned contrasts revealed trending differences between conditions for intergenerational dyads only, consistent with visual inspection (Figure 2) and descriptive statistics showing that state D is least represented when intergenerational dyads draw alone. In intergenerational dyads, state D indicates heightened within-brain connectivity in the younger dyad member.

We tentatively propose that the increased within-brain connectivity in a single dyad member might reflect reduced monitoring of the social environment (including the other dyad member) and/or heightened self-monitoring (Heggli et al., 2019; Mayo & Shamay-Tsoory, 2024). For example, intergenerational dyads draw alone may show less state D (relative to intergenerational dyads drawing together or same generation drawing in either condition), if the older dyad member engaged in heightened monitoring of the environment (i.e., the art at hand) and reduced monitoring of their behaviour relative to the younger dyad member. This aligns with existing findings that contralateral functional connectivity between IFG and TPJ is associated with higher-order social cognition, such as divergent thinking involved in creative drawing (Huang et al., 2024), regulating socially induced emotions (Grecucci et al., 2013), and controlling actions under averted gaze (Zillekens et al., 2019). Though the design of the present study is not sufficient to disentangle these potential mechanisms, future studies could incorporate additional measures to do so. These could include a measure of absorption in drawing, such as Rheinberg and colleagues’ Short Flow Scale (Engeser & Rheinberg, 2008; Rheinberg et al., 2023), which has items that capture absorption as well as social ease or worry during the drawing activity. The influence of direct and averted gaze could be captured using eye-tracking.

### 4.3 Asymmetric connectivity patterns during intergenerational collaboration and turn-taking dynamics

States F and G feature low-to-medium connectivity between brains and prominent within-brain connectivity between the bilateral IFG of one dyad member. State G, but not F, differed between real and pseudo dyads, specifically for intergenerational dyads drawing together. Here, the older dyad member shows marked within-brain connectivity. This pattern aligns with the asymmetric connectivity patterns characteristic of leader–follower dynamics described by Heggli and colleagues (2019, 2021), as well as Mayo and Shamay-Tsoory (2024). It also aligns with overt actor–observer dynamics observed using EEG by Li and colleagues (2025). Leader–follower and actor–observer dynamics are typically characterised by overt turn-taking (Heggli et al., 2019, 2021; Q. Li et al., 2025). Yet our behavioural findings suggest that longer durations of the asymmetrical state G are associated with reduced turn-taking behaviour, relative to shorter durations. The longer mean duration of state G may instead reflect subtle leader–follower roles during sustained mutual coordination, rather than the overt turn-taking in previous studies (Heggli et al., 2019, 2021; Q. Li et al., 2025). Subtle leader–follower roles during continuous mutual coordination can be explained by different predictive models for each dyad member (Heggli et al., 2019; Mayo & Shamay-Tsoory, 2024). For example, while drawing continuously and simultaneously, the leader plans their own behaviour, whereas the follower predicts and reacts to the leader’s behaviour (Mayo & Shamay-Tsoory, 2024). In this context, the heightened within-brain connectivity that we observed for the older dyad member may indicate a leader role during simultaneous drawing with higher reliance on self-models. Of note, our research group has also found that intergenerational dyads’ drawings were rated more collaborative than same generation dyads’ drawings based on the coherence of the motifs and the use of space (reference redacted for peer review), supporting our tentative proposal that leader–follower roles emerged more frequently in intergenerational than same generation dyads. Nonetheless, future research should include more refined measures of behaviour, such as the kinematics of turn-taking behaviour or gaze tracking, to better understand the dynamics of the social interaction associated with state G.

The opposite pattern, indicative of the older dyad member adapting to the younger leader (i.e., state F), did differ significantly between pseudo and real dyads in terms of coverage, showing that this state is also unique to real interaction. Yet, because the coverage of state F did not differ significantly between drawing tasks or groups, this state where younger dyad members lead and older dyad members follow is not informative as to the dynamics of intergenerational collaboration.

The asymmetrical pattern of within-brain connectivity involves prominently elevated connectivity between the bilateral IFG. The bilateral IFG are involved in language processing (Bulut, 2022), inhibitory control (Hampshire et al., 2010), attention reorientation (Frank & Sabatinelli, 2012), and social cognition (Diveica et al., 2021; Tso et al., 2018). Across these functions, unilateral left IFG activation is more common than unilateral right IFG activation (Jackson, 2021; Krieger-Redwood et al., 2015), though healthy older adults tend to show greater right IFG activation (Hoffman & Morcom, 2018). In the silent drawing task, explicit speech processing can be excluded as a possible mechanism for the observed connectivity between the bilateral IFG. Though implicit speech processing cannot be excluded, inhibitory control and/or attention reorientation seem to be more viable contributors. To establish the provenance of the bilateral IFG connectivity, additional measures of behaviour may also be useful to gain deeper insights into the pattern of within-brain connectivity observed in state G.

The strong effect of G in the intergenerational group might be additionally related to this finding. Building on these results, it seems reasonable to assume that states F and G reflect some semantic or inhibitory processes related to the task demands. Both inhibition of spontaneous impulses and semantic reasoning about the process or output can be speculated to be involved in drawing collaboratively.

### 4.4 Methodological considerations

#### 4.4.1 Age-related considerations

Age-related change in brain structure, cognition and functional connectivity are all highly interdependent and relevant considerations, each of which presents a potential confound for the interpretation of the results presented in this study. We accounted for age-related differences in haemodynamic signals stemming from differences in head and brain tissue structure using age-specific partial pathlength factors. However, the algorithm is currently only validated up to the age of 70 years (Scholkmann & Wolf, 2013). Validation of this algorithm for older adults would be valuable for replication of this work and for future intergenerational hyperscanning research conducted with fNIRS.

Age-related changes in cognition (Grady, 2012) and functional connectivity (Betzel et al., 2014; Grady, 2012; Pan et al., 2024) are open, and underdiscussed, challenges for the researchers conducting intergenerational hyperscanning studies (e.g., parent–child, student–teacher or younger–older adult). In this study, we overcome these potential confounds by comparing real and pseudo dyads (both same generation and intergenerational). In doing so, we can identify two-brain states where real dyads differ significantly from pseudo dyads in a metric (i.e., occurrence, duration, or coverage). If group differences in the identified states were driven by the inclusion of an older adult and not the direct social interaction between a younger and an older adult, real and pseudo intergenerational pairs would show comparable levels of the specific metric. This approach allows us to examine the processes underpinning collaborative social interactions between individuals with (potentially very) different cognitive abilities.

In this study, we observed that the duration of state G increased as turn-taking decreased for both groups. Zooming in on intergenerational dyads, we observed trends of lower state G durations for larger age differences and heightened age of the older dyad member, as well as parallel increases in overt turn-taking and the older dyad member’s age. We interpret these findings tentatively: It is plausible that during intergenerational collaboration, the older dyad member’s age shapes the amount of overt turn-taking during collaborative drawing, and plausibly by extension, the emergence and duration of state G. To shed light on the existence of any such causal relationship, carefully designed future studies may quantify age, cognitive abilities, social behaviours such as turn-taking, and two-brain states during intergenerational collaboration. We also note that differences in common ground, which can be amplified by differences in cumulative life experience (and lived experiences), may play an important role in collaborative behaviour, and potentially in two-brain states (Speer et al., 2024).

#### 4.4.2 State-based analysis considerations

Recently, there have been calls for developing data analysis techniques specific to fNIRS (Näher et al., 2024; Shin et al., 2024), as the scientific edge that fMRI and EEG approaches currently have over fNIRS might be partly due to the fact that a much richer and more mature range of methodologies specifically designed for the output of these methods already exists while fNIRS data still have yet to be fully exploited (Näher et al., 2024). Current computational models further emphasize the role of reconfigurations of neural dynamics within and between brains over the course of social interaction (Mayo & Shamay-Tsoory, 2024). The two-brain state analysis presented in this paper moves beyond averaging INS across tasks to a more holistic view that illuminates the temporal and spatial structure of two-brain dynamics, capturing the heterogeneity and asymmetry inherent in naturalistic interactions. The approach further extends previous approaches for characterizing two-brain states in fNIRS data (R. Li et al., 2021) as it integrates information on within-brain and between-brain connectivity and it goes beyond recently presented techniques for EEG data (Q. Li et al., 2025), since it can capture between-brain connectivity in addition to observing two interacting brains in parallel.

It should be noted that a state-based analysis is one way of summarising continuous, high-dimensional data that captures the dominant structure and relative differences. Further research will show whether the states generalise across individuals and studies. Importantly, dynamic functional connectivity often displays a poor signal-to-noise ratio and hence low reliability (Laumann et al., 2024). Future analyses could investigate further kernel functions (Wang et al., 2015), different window sizes, a meta-criterion for selecting the optimal clustering, as well as additional regions of interest. Regarding kernel functions, Näher and colleagues (2024) present analyses based on the super kernel matrix method where hyperparameter tuning happens separately for both chromophores which has been shown to lead to better classification results and might also be beneficial in clustering. Finally, a more flexible procedure could involve clustering on the dyad-level first and then identifying which states that replicate across dyads and relate to the behavioural structure of the experiment (Y. Zhang & Zhu, 2020). With respect to window lengths, we opted for one previously applied to fNIRS data. Longer windows may be beneficial for fNIRS data to capture slower oscillations relating to behaviour (Hakim et al., 2023). Future research may also examine the methodological strengths of aggregating criteria for selecting the optimal clustering into one meta-criterion (Michel & Koenig, 2018).

## 5 Conclusion

This study demonstrates that two-brain states can be identified in fNIRS hyperscanning signals collected in a naturalistic setting, such as a collaborative drawing task. We identified seven distinct two-brain states that arise during social interactions, three of which are specific to real interactions. Of these, six showed no clear evidence, and at most trends, for differing temporal dynamics per dyad composition or drawing condition. A single two-brain state (featuring low-to-medium connectivity between brains and prominent within-brain connectivity bilateral IFG of one dyad member) was observed for longer periods of time in intergenerational dyads than same generation dyads, during collaborative drawing. The duration of this state was negatively associated with turn-taking behaviour. None of the identified two-brain states showed evidence of longitudinal changes. Although only a subset of identified brain states showed meaningful relationships with the collaboration context, these findings substantiate the claim that asymmetric neural dynamics play a role during social interaction and depend on alternating patterns of within-brain and between-brain connectivity. A closer examination of such reconfigurations in naturalistic social interaction necessitates approaches that preserve spatial and temporal information, supporting the utility of this and similar methods for future work.

## Acknowledgments

We thank the survey center operated by UZH Healthy Longevity Center, University of Zurich, for their assistance with recruitment of older adults from the Zurich community. We thank Tessa Portier and Medea Häuselmann for their assistance with data collection, and Linda Fanconi for scheduling sessions. We thank Filiz Kanele for proofreading.

## Glossary

*Brain state*: A recurrent spatial pattern of brain activity
*Common cognitive processing*: The theory that similar perception of stimuli causes interacting people’s brains to respond in comparable ways at comparable time delays to the stimuli
*Coverage*: A brain state metric that indexes prevalence; the total number of windows in which a given brain state is active
*Duration*: A brain state metric that indexes state stability; the mean number of consecutive windows in which a given brain state is identified
*Hyperparameter tuning*: Selection of appropriate values for model parameters that are not learned from the data itself (e.g., shrinkage parameter or number of clusters) by evaluating model performance across a range of values (the hyperparameter space) and choosing the one that best meets a predefined criterion
*Hyperscanning*: Simultaneous recording of brain activity from two or more people while they interact
*Kernel function*: A (mathematical) function that quantifies similarity between two observations in a feature space without explicitly computing their coordinates in that space
*Kernel matrix*: A square matrix containing kernel function values for all pairs of observations, quantifying their mutual similarities
*Mutual prediction*: The theory that people continuously predict potential changes in their shared environments, including social information about each other
*Occurrence*: A brain state metric that indexes frequency; the number of discrete entries into a given brain state

## References

Aron, A., Aron, E. N., & Smollan, D. (1992). Inclusion of other in the self scale and the structure of interpersonal closeness. Journal of Personality and Social Psychology, 63(4), 17. 10.1037/0022-3514.63.4.596

Barachant, A., Quentin Barthélemy, Gabriel Wagner vom Berg, Alexandre Gramfort, Jean-Rémi King, Pedro L. C. Rodrigues, Bruna Junqueira Lopes, gcattan, Dave, Emanuele Olivetti, Vladislav Goncharenko, Maxence Dolle, toncho11, Tim Näher, Maria Sayu Yamamoto, GhilesReguig Apolline Mellot, Ammar Mian stonebig, … Frida Heskebeck. (2026). pyRiemann/pyRiemann: V0.10 (Version v0.10) [Computer software]. Zenodo. 10.5281/ZENODO.593816

Bates, D., Mächler, M., Bolker, B., & Walker, S. (2015). Fitting linear mixed-effects models using lme4. Journal of Statistical Software, 67(1), 1–48. 10.18637/jss.v067.i01

Benjamini, Y., & Hochberg, Y. (1995). Controlling the false discovery rate: A practical and powerful approach to multiple testing. Journal of the Royal Statistical Society: Series B (Methodological), 57(1), 289–300. 10.1111/j.2517-6161.1995.tb02031.x

Betzel, R. F., Byrge, L., He, Y., Goñi, J., Zuo, X.-N., & Sporns, O. (2014). Changes in structural and functional connectivity among resting-state networks across the human lifespan. NeuroImage, 102, 345–357. 10.1016/j.neuroimage.2014.07.067

Bilek, E., Zeidman, P., Kirsch, P., Tost, H., Meyer-Lindenberg, A., & Friston, K. (2022). Directed coupling in multi-brain networks underlies generalized synchrony during social exchange. NeuroImage, 252, 119038. 10.1016/j.neuroimage.2022.119038

Bollen, K. A., & Jackman, R. W. (1990). Regression Diagnostics: An Expository Treatment of Outliers and Influential Cases in Modern Methods of Data Analysis. In J. Fox & J. S. Long (Eds.), Modern methods of data analysis. Sage Publications.

Brigadoi, S., & Cooper, R. J. (2015). How short is short? Optimum source–detector distance for short-separation channels in functional near-infrared spectroscopy. Neurophotonics, 2(2), 025005. 10.1117/1.nph.2.2.025005

Bulut, T. (2022). Meta-analytic connectivity modeling of the left and right inferior frontal gyri. Cortex, 155, 107–131. 10.1016/j.cortex.2022.07.003

Burgess, A. P. (2013). On the interpretation of synchronization in EEG hyperscanning studies: A cautionary note. Frontiers in Human Neuroscience, 7. 10.3389/fnhum.2013.00881

Calinski, T., & Harabasz, J. (1974). A dendrite method for cluster analysis. Communications in Statistics - Theory and Methods, 3(1), 1–27. 10.1080/03610927408827101

Cross, E. S., Kraemer, D. J. M., Hamilton, A. F. D. C., Kelley, W. M., & Grafton, S. T. (2009). Sensitivity of the Action Observation Network to Physical and Observational Learning. Cerebral Cortex, 19(2), 315–326. 10.1093/cercor/bhn083

Czeszumski, A., Eustergerling, S., Lang, A., Menrath, D., Gerstenberger, M., Schuberth, S., Schreiber, F., Rendon, Z. Z., & König, P. (2020). Hyperscanning: A valid method to study neural inter-brain underpinnings of social interaction. Frontiers in Human Neuroscience, 14, 39. 10.3389/fnhum.2020.00039

Czeszumski, A., Liang, S. H.-Y., Dikker, S., König, P., Lee, C.-P., Koole, S. L., & Kelsen, B. (2022). Cooperative behavior evokes interbrain synchrony in the prefrontal and temporoparietal cortex: A systematic review and meta-analysis of fNIRS hyperscanning studies. eNeuro, 9(2), ENEURO.0268-21.2022. 10.1523/ENEURO.0268-21.2022

Dai, B., Zhai, Y., Long, Y., & Lu, C. (2024). How the Listener’s Attention Dynamically Switches Between Different Speakers During a Natural Conversation. Psychological Science, 35(6), 635–652. 10.1177/09567976241243367

Davies, D. L., & Bouldin, D. W. (1979). A Cluster Separation Measure. IEEE Transactions on Pattern Analysis and Machine Intelligence, PAMI-1(2), 224–227. 10.1109/TPAMI.1979.4766909

De Felice, S., Chand, T., Croy, I., Engert, V., Goldstein, P., Holroyd, C. B., Kirsch, P., Krach, S., Ma, Y., Scheele, D., Schurz, M., Schweinberger, S. R., Hoehl, S., & Vrticka, P. (2025). Relational neuroscience: Insights from hyperscanning research. Neuroscience & Biobehavioral Reviews, 169, 105979. 10.1016/j.neubiorev.2024.105979

De Felice, S., Hakim, U., Gunasekara, N., Pinti, P., Tachtsidis, I., & Hamilton, A. (2024). Having a chat and then watching a movie: How social interaction synchronises our brains during co-watching. Oxford Open Neuroscience, 3, kvae006. 10.1093/oons/kvae006

De Jong Gierveld, J., & van Tilburg, T. (2006). A 6-item scale for overall, emotional, and social loneliness: Confirmatory tests on survey data. Research on Aging, 28(5), 582–598. 10.1177/0164027506289723

DePasquale, C. E. (2020). A systematic review of caregiver–child physiological synchrony across systems: Associations with behavior and child functioning. Development and Psychopathology, 32(5), 1754–1777. 10.1017/S0954579420001236

Dikker, S., Michalareas, G., Oostrik, M., Serafimaki, A., Kahraman, H. M., Struiksma, M. E., & Poeppel, D. (2021). Crowdsourcing neuroscience: Inter-brain coupling during face-to-face interactions outside the laboratory. NeuroImage, 227, 117436. 10.1016/j.neuroimage.2020.117436

Dikker, S., Montgomery, S., & Tunca, T. (2019). Using synchrony-based neurofeedback in search of human connectedness. In Brain Art: Brain-Computer Interfaces for Artistic Expression (pp. 161–206). Springer.

Diveica, V., Koldewyn, K., & Binney, R. J. (2021). Establishing a role of the semantic control network in social cognitive processing: A meta-analysis of functional neuroimaging studies. NeuroImage, 245, 118702. 10.1016/j.neuroimage.2021.118702

Engeser, S., & Rheinberg, F. (2008). Flow, performance and moderators of challenge-skill balance. Motivation and Emotion, 32(3), 158–172. 10.1007/s11031-008-9102-4

Finn, E. S., Glerean, E., Khojandi, A. Y., Nielson, D., Molfese, P. J., Handwerker, D. A., & Bandettini, P. A. (2020). Idiosynchrony: From shared responses to individual differences during naturalistic neuroimaging. NeuroImage, 215, 116828. 10.1016/j.neuroimage.2020.116828

Fishburn, F. A., Ludlum, R. S., Vaidya, C. J., & Medvedev, A. V. (2019). Temporal Derivative Distribution Repair (TDDR): A motion correction method for fNIRS. NeuroImage, 184, 171–179. 10.1016/j.neuroimage.2018.09.025

Frank, D. W., & Sabatinelli, D. (2012). Stimulus-driven reorienting in the ventral frontoparietal attention network: The role of emotional content. Frontiers in Human Neuroscience, 6. 10.3389/fnhum.2012.00116

Garcia-Castro, G. (n.d.). Implementing the Differential Pathlength Factor (DPF) in Python: The Scholkmann Method. Retrieved https://gongcastro.github.io/blog/dpf-scholkmann/dpf-scholkmann.html

Garnier, S., Ross, N., BoB Rudis, Filipovic-Pierucci, A., Galili, T. Timelyportfolio, O’Callaghan, A., Greenwell, B., Sievert, C., Harris, D. J., Sciaini, M., & JJ Chen. (2023). sjmgarnier/viridis: CRAN release v0.6.3 (Version v0.6.3CRAN) [Computer software]. Zenodo. 10.5281/ZENODO.4679423

Grady, C. (2012). The cognitive neuroscience of ageing. Nature Reviews Neuroscience, 13(7), 491–505. 10.1038/nrn3256

Gramfort, A., Luessi, M., Larson, E., Engemann, D. A., Strohmeier, D., Brodbeck, C., Parkkonen, L., & Hämäläinen, M. S. (2014). MNE software for processing MEG and EEG data. NeuroImage, 86, 446–460. 10.1016/j.neuroimage.2013.10.027

Grecucci, A., Giorgetta, C., Bonini, N., & Sanfey, A. G. (2013). Reappraising social emotions: The role of inferior frontal gyrus, temporo-parietal junction and insula in interpersonal emotion regulation. Frontiers in Human Neuroscience, 7. 10.3389/fnhum.2013.00523

Gvirts, H., Ehrenfeld, L., Sharma, M., & Mizrahi, M. (2023). Virtual social interactions during the COVID-19 pandemic: The effect of interpersonal motor synchrony on social interactions in the virtual space. Scientific Reports, 13(1), 10481. 10.1038/s41598-023-37218-6

Hakim, U., De Felice, S., Pinti, P., Zhang, X., Noah, J., Ono, Y., Burgess, P. W., Hamilton, A., Hirsch, J., & Tachtsidis, I. (2023). Quantification of inter-brain coupling: A review of current methods used in haemodynamic and electrophysiological hyperscanning studies. NeuroImage, 280, 120354. 10.1016/j.neuroimage.2023.120354

Hamilton, A. F. de C. (2021). Hyperscanning: Beyond the hype. Neuron, 109(3), 404–407. 10.1016/j.neuron.2020.11.008

Hampshire, A., Chamberlain, S. R., Monti, M. M., Duncan, J., & Owen, A. M. (2010). The role of the right inferior frontal gyrus: Inhibition and attentional control. NeuroImage, 50(3), 1313–1319. 10.1016/j.neuroimage.2009.12.109

Happé, F., Cook, J. L., & Bird, G. (2017). The Structure of Social Cognition: In(ter)dependence of Sociocognitive Processes. Annual Review of Psychology, 68(1), 243–267. 10.1146/annurev-psych-010416-044046

Hasson, U., Ghazanfar, A. A., Galantucci, B., Garrod, S., & Keysers, C. (2012). Brain-to-brain coupling: A mechanism for creating and sharing a social world. Trends in Cognitive Sciences, 16(2), 114–121. 10.1016/j.tics.2011.12.007

Heggli, O. A., Cabral, J., Konvalinka, I., Vuust, P., & Kringelbach, M. L. (2019). A Kuramoto model of self-other integration across interpersonal synchronization strategies. PLOS Computational Biology, 15(10), e1007422. 10.1371/journal.pcbi.1007422

Heggli, O. A., Konvalinka, I., Cabral, J., Brattico, E., Kringelbach, M. L., & Vuust, P. (2021). Transient brain networks underlying interpersonal strategies during synchronized action. Social Cognitive and Affective Neuroscience, 16(1– 2), 19–30. 10.1093/scan/nsaa056

Hendrikse, S. C. F., Treur, J., Wilderjans, T. F., Dikker, S., & Koole, S. L. (2025). The Effect of Transition Dynamics in Synchrony on Social Interaction Adaptivity: A Multi-adaptive Network Model. In S. C. F. Hendrikse, J. Treur, & S. L. Koole (Eds.), New Analysis and Modeling Directions in Social Interaction Science (Vol. 614, pp. 465–493). Springer Nature Switzerland. 10.1007/978-3-031-99968-0_15

Hoehl, S., Fairhurst, M., & Schirmer, A. (2021). Interactional synchrony: Signals, mechanisms and benefits. Social Cognitive and Affective Neuroscience, 16(1–2), 5–18. 10.1093/scan/nsaa024

Hoffman, P., & Morcom, A. M. (2018). Age-related changes in the neural networks supporting semantic cognition: A meta-analysis of 47 functional neuroimaging studies. Neuroscience & Biobehavioral Reviews, 84, 134–150. 10.1016/j.neubiorev.2017.11.010

Huang, F., Fu, X., Song, J., Ren, J., Li, F., & Zhao, Q. (2024). Divergent thinking benefits from functional antagonism of the left IFG and right TPJ: A transcranial direct current stimulation study. Cerebral Cortex, 34(2), bhad531. 10.1093/cercor/bhad531

Jackson, R. L. (2021). The neural correlates of semantic control revisited. NeuroImage, 224, 117444. 10.1016/j.neuroimage.2020.117444

Kingsbury, L., Huang, S., Wang, J., Gu, K., Golshani, P., Wu, Y. E., & Hong, W. (2019). Correlated neural activity and encoding of behavior across brains of socially interacting animals. Cell, 178(2), 429–446.e16. 10.1016/j.cell.2019.05.022

Krieger-Redwood, K., Teige, C., Davey, J., Hymers, M., & Jefferies, E. (2015). Conceptual control across modalities: Graded specialisation for pictures and words in inferior frontal and posterior temporal cortex. Neuropsychologia, 76, 92–107. 10.1016/j.neuropsychologia.2015.02.030

Laumann, T. O., Snyder, A. Z., & Gratton, C. (2024). Challenges in the measurement and interpretation of dynamic functional connectivity. Imaging Neuroscience, 2, imag–2–00366. 10.1162/imag_a_00366

Lenth, R. V. (2021). emmeans: Estimated marginal means, aka least-squares means [Computer software].

Li, Q., Zimmermann, M., & Konvalinka, I. (2025). Two-brain microstates: A novel hyperscanning-EEG method for quantifying task-driven inter-brain asymmetry. Journal of Neuroscience Methods, 416, 110355. 10.1016/j.jneumeth.2024.110355

Li, R., Mayseless, N., Balters, S., & Reiss, A. L. (2021). Dynamic inter-brain synchrony in real-life inter-personal cooperation: A functional near-infrared spectroscopy hyperscanning study. NeuroImage, 238, 118263. 10.1016/j.neuroimage.2021.118263

Lu, J., Wang, Y., Shu, Z., Zhang, X., Wang, J., Cheng, Y., Zhu, Z., Yu, Y., Wu, J., Han, J., & Yu, N. (2022). fNIRS-based brain state transition features to signify functional degeneration after Parkinson’s disease. Journal of Neural Engineering, 19(4), 046038. 10.1088/1741-2552/ac861e

Luke, R., Larson, E., Shader, M. J., Innes-Brown, H., Van Yper, L., Lee, A. K. C., Sowman, P. F., & McAlpine, D. (2021). Analysis methods for measuring passive auditory fNIRS responses generated by a block-design paradigm. Neurophotonics, 8(02). 10.1117/1.NPh.8.2.025008

Marton-Alper, I. Z., Markus, A., Nevat, M., Bennet, R., & Shamay-Tsoory, S. G. (2023). Differential contribution of between and within-brain coupling to movement synchronization. Human Brain Mapping, 44(10), 4136–4151. 10.1002/hbm.26335

Mayo, O., & Gordon, I. (2020). In and out of synchrony—Behavioral and physiological dynamics of dyadic interpersonal coordination. Psychophysiology, 57(6), e13574. 10.1111/psyp.13574

Mayo, O., & Shamay-Tsoory, S. (2024). Dynamic mutual predictions during social learning: A computational and interbrain model. Neuroscience & Biobehavioral Reviews, 157, 105513. 10.1016/j.neubiorev.2023.105513

Michel, C. M., & Koenig, T. (2018). EEG microstates as a tool for studying the temporal dynamics of whole-brain neuronal networks: A review. NeuroImage, 180, 577–593. 10.1016/j.neuroimage.2017.11.062

Moffat, R. (2024). Invisible mechanisms of interpersonal alignment. Nature Reviews Psychology. 10.1038/s44159-024-00284-2

Munafò, M. R., Nosek, B. A., Bishop, D. V. M., Button, K. S., Chambers, C. D., Percie Du Sert, N., Simonsohn, U., Wagenmakers, E.-J., Ware, J. J., & Ioannidis, J. P. A. (2017). A manifesto for reproducible science. Nature Human Behaviour, 1(1), 0021. 10.1038/s41562-016-0021

Näher, T., Bastian, L., Vorreuther, A., Fries, P., Goebel, R., & Sorger, B. (2024). Riemannian Geometry for the classification of brain states with fNIRS. Neuroscience. 10.1101/2024.09.06.611347

Oostenveld, R., Fries, P., Maris, E., & Schoffelen, J.-M. (2011). Fieldtrip: Open source software for advanced analysis of MEG, EEG, and invasive electrophysiological data. Computational Intelligence and Neuroscience, 2011, 1–9. 10.1155/2011/156869

Pan, Y., Bi, C., Kochunov, P., Shardell, M., Smith, J. C., McCoy, R. G., Ye, Z., Yu, J., Lu, T., Yang, Y., Lee, H., Liu, S., Gao, S., Ma, Y., Li, Y., Chen, C., Ma, T., Wang, Z., Nichols, T., … Chen, S. (2024). Brain-wide functional connectome analysis of 40,000 individuals reveals brain networks that show aging effects in older adults. Imaging Neuroscience, 2, imag–2–00394. 10.1162/imag_a_00394

Park, H.-J., & Friston, K. (2013). Structural and Functional Brain Networks: From Connections to Cognition. Science, 342(6158), 1238411. 10.1126/science.1238411

Paulus, C. (2009). Der Saarbrücker Persönlichkeitsfrageboden SPF (IRI) zur Messung von Empathie: Psychometrische Evaluation der deutschen Version des Interpersonal Reactivity Index. 10.23668/psycharchives.9249

Pauly, T., Gerstorf, D., Wahl, H.-W., & Hoppmann, C. A. (2021). A developmental–contextual model of couple synchrony across adulthood and old age. Psychology and Aging, 36(8), 943–956. 10.1037/pag0000651

Pedregosa, F., Varoquaux, G., Gramfort, A., Michel, V., Thirion, B., Grisel, O., Blondel, M., Prettenhofer, P., Weiss, R., Dubourg, V., Vanderplas, J., Passos, A., Cournapeau, D., Brucher, M., Perrot, M., & Duchesnay, E. (2011). Scikit-learn: Machine Learning in Python. 12, 2825–2830.

Peirce, J. W., Hirst, R. J., & MacAskill, M. R. (2022). Building Experiments in PsychoPy (2nd ed.). Sage. https://uk.sagepub.com/en-gb/eur/building-experiments-in-psychopy/book273700

Pittinsky, T. L., Rosenthal, S. A., & Montoya, R. M. (2011). Measuring positive attitudes toward outgroups: Development and validation of the Allophilia Scale. In L. R. Tropp & R. K. Mallett (Eds.), Moving beyond prejudice reduction: Pathways to positive intergroup relations. (pp. 41–60). American Psychological Association. 10.1037/12319-002

Pollonini, L., Olds, C., Abaya, H., Bortfeld, H., Beauchamp, M. S., & Oghalai, J. S. (2014). Auditory cortex activation to natural speech and simulated cochlear implant speech measured with functional near-infrared spectroscopy. Hearing Research, 309, 84–93. 10.1016/j.heares.2013.11.007

Rheinberg, F., Vollmeyer, R., Engster, S., & Sreeramoju, R. R. (2023). FSS – Flow short scale (English version). In J. Stiensmeier-Pelster & F. Rheinberg (Eds.), The Acquisition of the Flow Experience (pp. 261–279).

Roche, E. C., Redcay, E., & Romeo, R. R. (2025). Caregiver-child neural synchrony: Magic, mirage, or developmental mechanism? Developmental Cognitive Neuroscience, 71, 101482. 10.1016/j.dcn.2024.101482

Schilbach, L., Timmermans, B., Reddy, V., Costall, A., Bente, G., Schlicht, T., & Vogeley, K. (2013). Toward a second-person neuroscience. Behavioral and Brain Sciences, 36(4), 393–414. 10.1017/S0140525X12000660

Scholkmann, F., & Wolf, M. (2013). General equation for the differential pathlength factor of the frontal human head depending on wavelength and age. Journal of Biomedical Optics, 18(10), 105004. 10.1117/1.JBO.18.10.105004

Schurz, M., Radua, J., Tholen, M. G., Maliske, L., Margulies, D. S., Mars, R. B., Sallet, J., & Kanske, P. (2021). Toward a hierarchical model of social cognition: A neuroimaging meta-analysis and integrative review of empathy and theory of mind. Psychological Bulletin, 147(3), 293–327. 10.1037/bul0000303

Shamay-Tsoory, S. G. (2022). Brains that Fire Together Wire Together: Interbrain Plasticity Underlies Learning in Social Interactions. The Neuroscientist, 28(6), 543–551. 10.1177/1073858421996682

Shamay-Tsoory, S. G., Saporta, N., Marton-Alper, I. Z., & Gvirts, H. Z. (2019). Herding brains: A core neural mechanism for social alignment. Trends in Cognitive Sciences, 23(3), 174–186. 10.1016/j.tics.2019.01.002

Shawe-Taylor, J., & Cristianini, N. (2004). Kernel Methods for Pattern Analysis. Cambridge University Press.

Shin, J. H., Kang, M. J., & Lee, S. A. (2024). Wearable functional near-infrared spectroscopy for measuring dissociable activation dynamics of prefrontal cortex subregions during working memory. Human Brain Mapping, 45(2), e26619. 10.1002/hbm.26619

Speer, S. P. H., Mwilambwe-Tshilobo, L., Tsoi, L., Burns, S. M., Falk, E. B., & Tamir, D. I. (2024). Hyperscanning shows friends explore and strangers converge in conversation. Nature Communications, 15(1), 7781. 10.1038/s41467-024-51990-7

Tso, I. F., Rutherford, S., Fang, Y., Angstadt, M., & Taylor, S. F. (2018). The “social brain” is highly sensitive to the mere presence of social information: An automated meta-analysis and an independent study. PLOS ONE, 13(5), e0196503. 10.1371/journal.pone.0196503

United Nations. (2024). World population prospects 2024: Summary of results. United Nations.

Venables, W. N., & Ripley, B. D. (2002). Modern applied statistics with S (4th ed). Springer.

Wang, L., Zhang, J., Zhou, L., Tang, C., & Li, W. (2015). Beyond Covariance: Feature Representation with Nonlinear Kernel Matrices. 2015 IEEE International Conference on Computer Vision (ICCV), 4570–4578. 10.1109/ICCV.2015.519

Wickham, H. (2016). ggplot2: Elegant Graphics for Data Analysis (2nd ed. 2016). Springer International Publishing : Imprint: Springer. 10.1007/978-3-319-24277-4

Wickham, H., Averick, M., Bryan, J., Chang, W., McGowan, L., François, R., Grolemund, G., Hayes, A., Henry, L., Hester, J., Kuhn, M., Pedersen, T., Miller, E., Bache, S., Müller, K., Ooms, J., Robinson, D., Seidel, D., Spinu, V., … Yutani, H. (2019). Welcome to the Tidyverse. Journal of Open Source Software, 4(43), 1686. 10.21105/joss.01686

Wickham, H., & Henry, L. (2026). purrr: Functional Programming Tools [Computer software].

Yücel, M. A., Lühmann, A. v., Scholkmann, F., Gervain, J., Dan, I., Ayaz, H., Boas, D., Cooper, R. J., Culver, J., Elwell, C. E., Eggebrecht, A., Franceschini, M. A., Grova, C., Homae, F., Lesage, F., Obrig, H., Tachtsidis, I., Tak, S., Tong, Y.,… Wolf, M. (2021). Best practices for fNIRS publications. Neurophotonics, 8(1), 1–34. 10.1117/1.nph.8.1.012101

Zeileis, A., & Hothorn, T. (2002). Diagnostic Checking in Regression Relationships (Pt. 3). 2, 7–10.

Zhang, L., Xu, X., Li, Z., Chen, L., & Feng, L. (2022). Interpersonal neural synchronization predicting learning outcomes from teaching-learning interaction: A meta-analysis. Frontiers in Psychology, 13, 835147. 10.3389/fpsyg.2022.835147

Zhang, Y., & Zhu, C. (2020). Assessing Brain Networks by Resting-State Dynamic Functional Connectivity: An fNIRS-EEG Study. Frontiers in Neuroscience, 13, 1430. 10.3389/fnins.2019.01430

Zillekens, I. C., Schliephake, L. M., Brandi, M.-L., & Schilbach, L. (2019). A look at actions: Direct gaze modulates functional connectivity of the right TPJ with an action control network. Social Cognitive and Affective Neuroscience, 14(9), 977–986. 10.1093/scan/nsz071

