## Supplemental Material for "Two-brain states characterise within- and between-brain connectivity during intergenerational collaborative drawing"

**Supplementary Materials:** [redacted for peer review]. *Two-brain states during collaborative drawing reflect leader–follower dynamics in intergenerational dyads.*


| Item | Pages | Item | Pages |
|---|---|---|---|
| ***Methods Details*** | | Supplementary Table 28 | 24 |
| Kernel functions | 2 | Supplementary Table 29 | 24 |
| Model selection | 2 | Supplementary Table 30 | 25 |
| | | Supplementary Table 31 | 25 |
| ***Figures*** | | Supplementary Table 32 | 26 |
| Supplementary Figure 1 | 3 | Supplementary Table 33 | 27 |
| Supplementary Figure 2 | 4 | Supplementary Table 34 | 28 |
| | | Supplementary Table 35 | 29 |
| ***Tables*** | | Supplementary Table 36 | 30 |
| Supplementary Table 1 | 5 | Supplementary Table 37 | 31 |
| Supplementary Table 2 | 5 | Supplementary Table 38 | 31 |
| Supplementary Table 3 | 6 | Supplementary Table 39 | 32 |
| Supplementary Table 4 | 7 | Supplementary Table 40 | 32 |
| Supplementary Table 5 | 8 | | |
| Supplementary Table 6 | 8 | ***Three-cluster solution*** | 33 |
| Supplementary Table 7 | 9 | | |
| Supplementary Table 8 | 10 | ***Shared brain states*** | 34 |
| Supplementary Table 9 | 11 | | |
| Supplementary Table 10 | 11 | ***References*** | 35 |
| Supplementary Table 11 | 12 | | |
| Supplementary Table 12 | 13 | | |
| Supplementary Table 13 | 14 | | |
| Supplementary Table 14 | 14 | | |
| Supplementary Table 15 | 15 | | |
| Supplementary Table 16 | 15 | | |
| Supplementary Table 17 | 16 | | |
| Supplementary Table 18 | 17 | | |
| Supplementary Table 19 | 18 | | |
| Supplementary Table 20 | 19 | | |
| Supplementary Table 21 | 20 | | |
| Supplementary Table 22 | 20 | | |
| Supplementary Table 23 | 21 | | |
| Supplementary Table 24 | 22 | | |
| Supplementary Table 25 | 22 | | |
| Supplementary Table 26 | 22 | | |
| Supplementary Table 27 | 23 | | |

**Supplementary Materials:** [redacted for peer review]. *Two-brain states during collaborative drawing reflect leader–follower dynamics in intergenerational dyads.*

**Supplementary Methods Information**

**Kernel functions.** We employed five different kernel functions: covariance, correlation coefficient, Tyler's M-estimator-based covariance, Ledoit–Wolf shrunk covariance, and the radial basis function. Covariance between two channel signals is defined as the expected value of the demeaned signals, capturing raw linear co-fluctuation. The correlation coefficient is the covariance normalised at the time series' standard deviations, rendering its values comparable across channels but discarding amplitude information. Tyler's M-estimator-based covariance is particularly useful when the data contains heavy-tailed distributions or outliers which would otherwise skew the covariance (Goes et al., 2020). The Ledoit–Wolf shrunk covariance stabilises covariance estimation by shrinking the sample covariance towards a well-conditioned target, trading robustness to outliers for estimation reliability (Ledoit & Wolf, 2004). The radial basis function is the exponential of the negative squared and scaled Euclidean distance between the two input time series, where the scaling parameter $\gamma$ was set to 1. It is a measure of non-linear similarity between both channel signals (Rocha, 2009). Together, these kernel functions span linear versus nonlinear dependence, scale-dependent versus scale-invariant coupling, and classical versus regularised covariance estimation.

**Model selection.** Since the outcome variables represent count or percentage data, we tested for each analysis whether a Poisson model would be appropriate or if overdispersion necessitates an approached based on the negative binomial distribution (Ver Hoef & Boveng, 2007). In all cases, the Poisson model exhibited overdispersion, violating the assumption that the mean equals the variance. Likelihood ratio tests to compare the AIC also significantly favoured the negative binomial model over the Poisson model, and therefore it was used in all analyses. Model comparisons are reported in Supplementary Table 3.

**Supplementary Materials:** [redacted for peer review]. *Two-brain states during collaborative drawing reflect leader–follower dynamics in intergenerational dyads.*

### Supplementary Figures

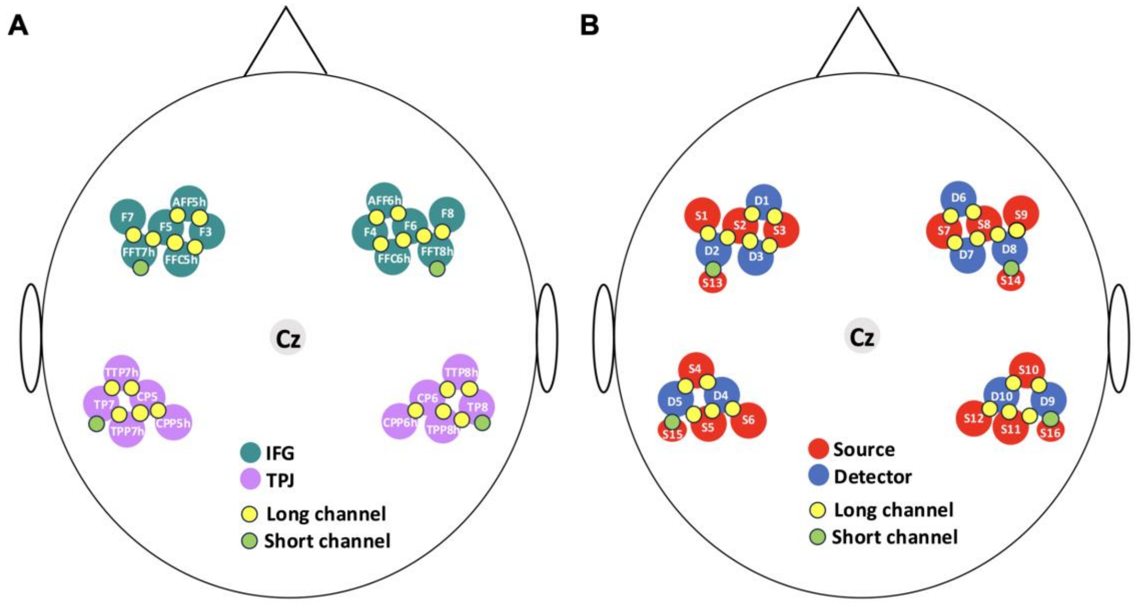


**Supplementary Figure 1.** A) EEG 5-10 positions of each optode within the four ROIs (bilateral IFG and TPJ). B) Arrangement optodes, where numbers correspond to sensor numbers on Cortivision Photon Cap Device (S13-S16 are sources specifically designed to measure short channel signals).

**Supplementary Materials:** [redacted for peer review]. *Two-brain states during collaborative drawing reflect leader–follower dynamics in intergenerational dyads.*

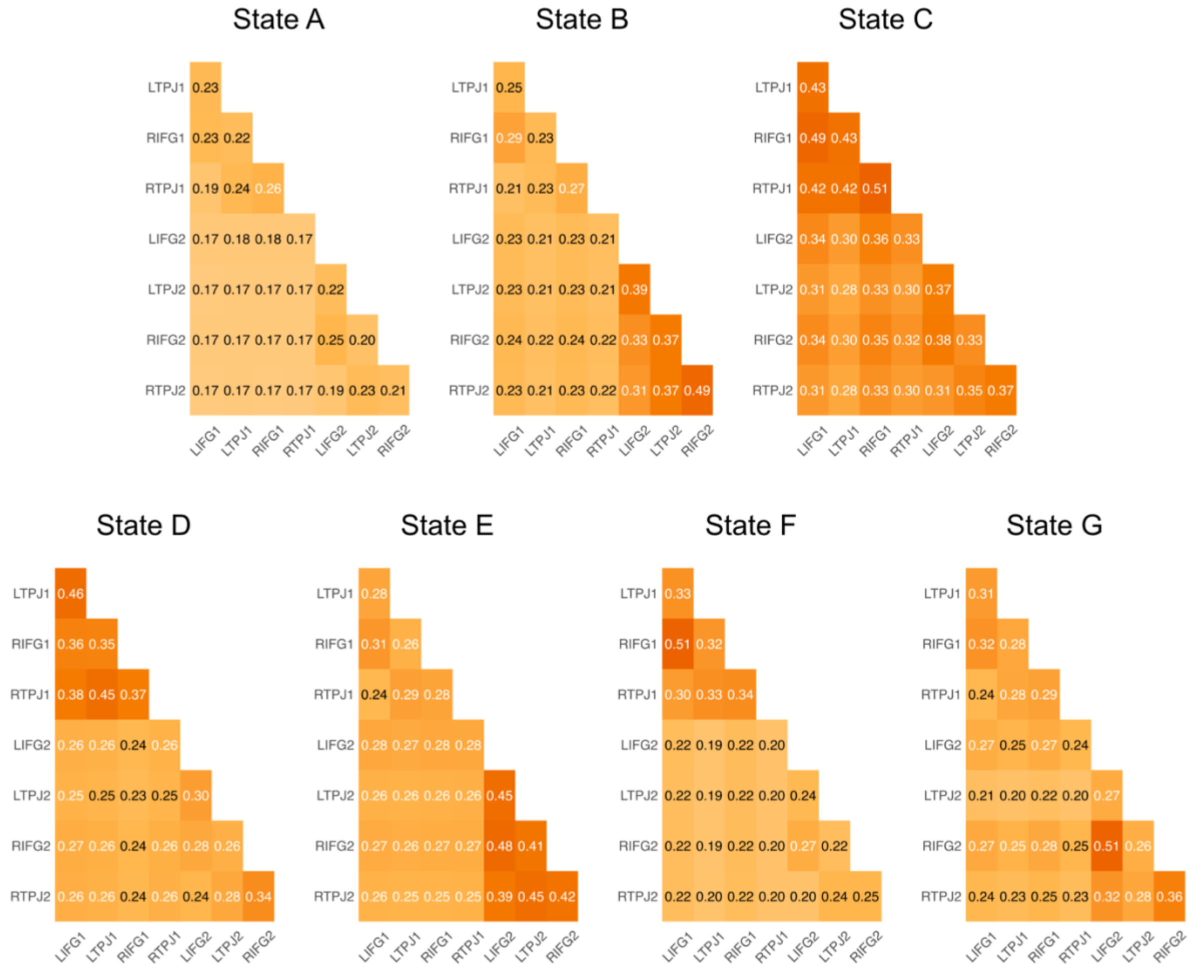


**Supplementary Figure 2.** Seven two-brain states identified using Riemannian geometry-based clustering of dynamic functional connectivity matrices. Matrices show similarities in the HbR signals between ROIs belonging to each dyad member (distinguished by 1 and 2).

**Supplementary Materials:** [redacted for peer review]. *Two-brain states during collaborative drawing reflect leader–follower dynamics in intergenerational dyads.*

### Supplementary Tables

**Supplementary Table 1.** Data exclusion. Breakdown of reasons for excluding dyads data. Recordings/dyads only included once in list. E.g., if a recording/dyad is counted among reasons unrelated to signal quality, they are not re-counted among reasons related to signal quality. In total 91 recordings of 732 recordings were excluded, impacting 85 of 366 dyadic sessions.

| Reason | N individual recordings excluded | N dyadic sessions impacted |
|---|---|---|
| ***Unrelated to signal quality*** | | |
| Recordings lost in transition between recording laptops | 2 | 2 |
| Triggers missing or unreliable | 7 | 4 |
| Participant unable to complete session (could not attend final session/consumed alcohol) | 5 | 3 |
| Multi-part recording (bluetooth or battery dropout) | 6 | 4 |
| ***Related to signal quality*** | | |
| No short channels of adequate quality | 71 | 69 |
| No long channels of adequate quality (pilot montage used) | 0 | 0 |
| **Total** | **91** | **85** |

**Supplementary Table 2.** Hyperparameter space.

| Hyperparameter | Search Space | Selected value (CHI) | Selected value (DBI) |
|---|---|---|---|
| Kernel function | {cov, corr, tyl, lwf, rbf} | rbf | rbf |
| Shrinkage parameter $\lambda$ | {0, 0.01, 0.1} | 0.1 | 0.1 |
| Number of clusters | {3, 4, 5, 6, 7, 8, 9, 10} | 3 | 7 |

*Note.* cov = covariance, corr = correlation coefficient, tyl = Tyler's M-estimator-based covariance, lwf = Ledoit–Wolf shrunk covariance, rbf = radial basis function, CHI = Calinski–Harabasz index, DBI = Davies–Bouldin index.

**Supplementary Materials:** [redacted for peer review]. *Two-brain states during collaborative drawing reflect leader–follower dynamics in intergenerational dyads.*

**Supplementary Table 3.** Model comparisons of Poisson and negative binomial models.

| **Metric** | **Model** | **Poisson model** | **Negative binomial model** | |
|---|---|---|---|---|
| | | **AIC** | **θ** | **AIC** |
| Occurrence | First step | 127502.4 | 8.65 | 124461.1 |
| Occurrence | Second step | 2830.7 | 5.70 | 2688.5 |
| Coverage | First step | 868952.0 | 1.40 | 248861.1 |
| Coverage | Second step | 28739.8 | 1.28 | 7457.6 |
| Duration | First step | 180846.9 | 4.26 | 153799.1 |
| Duration | Second step | 2032.4 | 2.98 | 1527.8 |

*Note.* First-step models refer to generalized linear models to test differences between real and pseudo dyads. Second-step models refer to models constructed only on data from real dyads and those states that differed between pseudo and real dyads in at least one condition in first-step models. AIC = Akaike Information Criterion.

**Supplementary Table 4.** Likelihood ratio test results with Chi-squared tests for occurrence, real and pseudo dyads.

| **Factor** | ***df*** | **Δ Deviance ($\chi^2$)** | ***p*** |
|---|---|---|---|
| Brain State | 6 | 2886.56 | < .001*** |
| Task | 1 | 2.28 | .131 |
| Dyad Type | 1 | 0.55 | .458 |
| Group | 1 | 0.01 | .924 |
| Session | 1 | 23.09 | < .001*** |
| Brain State × Task | 6 | 121.78 | < .001*** |
| Brain State × Dyad Type | 6 | 9.46 | .149 |
| Task × Dyad Type | 1 | 1.11 | .293 |
| Brain State × Group | 6 | 13.77 | .032* |
| Task × Group | 1 | 0.07 | .795 |
| Dyad Type × Group | 1 | 0.11 | .738 |
| Brain State × Session | 6 | 9.47 | .149 |
| Task × Session | 1 | 0.10 | .749 |
| Dyad Type × Session | 1 | 0.28 | .594 |
| Group × Session | 1 | 2.08 | .149 |
| Brain State × Task × Dyad Type | 6 | 2.08 | .912 |
| Brain State × Task × Group | 6 | 9.63 | .141 |
| Brain State × Dyad Type × Group | 6 | 17.69 | .007** |
| Task × Dyad Type × Group | 1 | 0.24 | .621 |
| Brain State × Task × Session | 6 | 7.38 | .287 |
| Brain State × Dyad Type × Session | 6 | 17.82 | .007** |
| Task × Dyad Type × Session | 1 | 1.01 | .314 |
| Brain State × Group × Session | 6 | 10.35 | .111 |
| Task × Group × Session | 1 | 0.51 | .476 |
| Dyad Type × Group × Session | 1 | 0.00 | .981 |
| Brain State × Task × Dyad Type × Group | 6 | 10.28 | .113 |
| Brain State × Task × Dyad Type × Session | 6 | 2.61 | .856 |
| Brain State × Task × Group × Session | 6 | 5.42 | .491 |
| Brain State × Dyad Type × Group × Session | 6 | 3.76 | .709 |
| Task × Dyad Type × Group × Session | 1 | 0.01 | .933 |
| Brain State × Task × Dyad Type × Group × Session | 6 | 12.61 | .0497* |

*Note.* Deviance tests are likelihood ratio $\chi^2$-tests from a negative binomial GLM (log link; θ = 8.65). The dependent variable is occurrence.

**Supplementary Table 5.** Contrast analyses for occurrence, real vs. pseudo dyads.

| Brain State | Task | Group | $z$ | $p_{adj}$ |
|---|---|---|---|---|
| A | Alone | Intergenerational | 0.850 | .837 |
| A | Alone | Same generation | 1.597 | .452 |
| A | Together | Intergenerational | 0.793 | .837 |
| A | Together | Same generation | 2.071 | .269 |
| B | Alone | Intergenerational | -0.446 | .837 |
| B | Alone | Same generation | 0.645 | .837 |
| B | Together | Intergenerational | -3.286 | .014* |
| B | Together | Same generation | -0.157 | .875 |
| C | Alone | Intergenerational | -0.448 | .837 |
| C | Alone | Same generation | -0.321 | .837 |
| C | Together | Intergenerational | 0.557 | .837 |
| C | Together | Same generation | -0.591 | .837 |
| D | Alone | Intergenerational | 3.385 | .014* |
| D | Alone | Same generation | -1.585 | .452 |
| D | Together | Intergenerational | 0.486 | .837 |
| D | Together | Same generation | -0.654 | .837 |
| E | Alone | Intergenerational | -1.315 | .660 |
| E | Alone | Same generation | -0.265 | .852 |
| E | Together | Intergenerational | -0.426 | .837 |
| E | Together | Same generation | 0.396 | .837 |
| F | Alone | Intergenerational | 0.942 | .837 |
| F | Alone | Same generation | 1.954 | .284 |
| F | Together | Intergenerational | 2.351 | .175 |
| F | Together | Same generation | -0.832 | .837 |
| G | Alone | Intergenerational | 0.394 | .837 |
| G | Alone | Same generation | -0.185 | .875 |
| G | Together | Intergenerational | -0.356 | .837 |
| G | Together | Same generation | -0.544 | .837 |

*Note*. *p*-values were adjusted for multiple comparisons using the false discovery rate (FDR).

**Supplementary Table 6.** Likelihood ratio test results with Chi-squared tests for occurrence, only real dyads.

| Factor | *df* | Δ Deviance ($\chi^2$) | *p* |
|---|---|---|---|
| Brain State | 1 | 6.25 | .012* |
| Task | 1 | 4.89 | .027* |
| Group | 1 | 0.36 | .549 |
| Brain State × Task | 1 | 0.00 | .984 |
| Brain State × Group | 1 | 3.20 | .074 |
| Task × Group | 1 | 1.39 | .238 |
| Brain State × Task × Group | 1 | 3.30 | .069 |

*Note.* Deviance tests are likelihood ratio $\chi^2$-tests from a negative binomial GLM (log link; $\theta = 5.70$). The dependent variable is occurrence in states B and D.

**Supplementary Table 7.** Contrast analyses for occurrence in states B and D, only real dyads.

| Contrast | State | $z$ | $p_{adj}$ |
|---|---|---|---|
| **Within-group task contrasts** | | | |
| Intergen: Alone vs. Together | B | -0.857 | .594 |
| Same gen: Alone vs. Together | B | -1.442 | .299 |
| Intergen: Alone vs. Together | D | -2.617 | .053 |
| Same gen: Alone vs. Together | D | 0.170 | .866 |
| **Within-task group contrasts** | | | |
| Alone: Intergen vs. Same gen | B | 0.726 | .624 |
| Together: Intergen vs. Same gen | B | 0.441 | .791 |
| Alone: Intergen vs. Same gen | D | -2.723 | .053 |
| Together: Intergen vs. Same gen | D | -0.250 | .865 |

*Note. p*-values were adjusted for multiple comparisons using the false discovery rate (FDR).

**Supplementary Table 8.** Likelihood ratio test results with Chi-squared tests for coverage, real and pseudo dyads.

| **Factor** | ***df*** | **Δ Deviance ($\chi^2$)** | ***p*** |
|---|---|---|---|
| Brain State | 6 | 1758.38 | < .001*** |
| Task | 1 | 4.84 | .028* |
| Dyad Type | 1 | 0.97 | .324 |
| Group | 1 | 0.00 | .962 |
| Session | 1 | 0.18 | .669 |
| Brain State × Task | 6 | 146.43 | < .001*** |
| Brain State × Dyad Type | 6 | 11.80 | .066 |
| Task × Dyad Type | 1 | 0.01 | .939 |
| Brain State × Group | 6 | 36.88 | < .001*** |
| Task × Group | 1 | 0.23 | .632 |
| Dyad Type × Group | 1 | 0.02 | .883 |
| Brain State × Session | 6 | 21.45 | .002** |
| Task × Session | 1 | 0.01 | .919 |
| Dyad Type × Session | 1 | 0.03 | .869 |
| Group × Session | 1 | 0.07 | .785 |
| Brain State × Task × Dyad Type | 6 | 1.43 | .964 |
| Brain State × Task × Group | 6 | 11.06 | .086 |
| Brain State × Dyad Type × Group | 6 | 28.99 | < .001*** |
| Task × Dyad Type × Group | 1 | 0.00 | .962 |
| Brain State × Task × Session | 6 | 3.32 | .768 |
| Brain State × Dyad Type × Session | 6 | 26.66 | < .001*** |
| Task × Dyad Type × Session | 1 | 0.00 | .993 |
| Brain State × Group × Session | 6 | 22.02 | .001** |
| Task × Group × Session | 1 | 0.01 | .940 |
| Dyad Type × Group × Session | 1 | 0.11 | .744 |
| Brain State × Task × Dyad Type × Group | 6 | 18.21 | .006** |
| Brain State × Task × Dyad Type × Session | 6 | 3.00 | .809 |
| Brain State × Task × Group × Session | 6 | 5.53 | .478 |
| Brain State × Dyad Type × Group × Session | 6 | 6.44 | .375 |
| Task × Dyad Type × Group × Session | 1 | 0.02 | .884 |
| Brain State × Task × Dyad Type × Group × Session | 6 | 1.99 | .921 |

*Note.* Deviance tests are likelihood ratio $\chi^2$-tests from a negative binomial GLM (log link; $\theta = 1.40$). The dependent variable is occurrence.

**Supplementary Table 9.** Contrast analyses for coverage, real vs. pseudo dyads.

| Brain State | Task | Group | $z$ | $p_{adj}$ |
|---|---|---|---|---|
| A | Alone | Intergenerational | 0.256 | .949 |
| A | Alone | Same generation | 1.373 | .432 |
| A | Together | Intergenerational | 0.544 | .863 |
| A | Together | Same generation | 2.217 | .124 |
| B | Alone | Intergenerational | -0.189 | .949 |
| B | Alone | Same generation | -0.891 | .652 |
| B | Together | Intergenerational | -2.978 | .027* |
| B | Together | Same generation | -0.631 | .821 |
| C | Alone | Intergenerational | -1.084 | .557 |
| C | Alone | Same generation | -1.131 | .556 |
| C | Together | Intergenerational | 1.627 | .325 |
| C | Together | Same generation | -1.773 | .305 |
| D | Alone | Intergenerational | 3.185 | .020* |
| D | Alone | Same generation | -1.623 | .325 |
| D | Together | Intergenerational | 1.311 | .442 |
| D | Together | Same generation | -0.678 | .820 |
| E | Alone | Intergenerational | -2.335 | .110 |
| E | Alone | Same generation | 0.300 | .949 |
| E | Together | Intergenerational | 0.307 | .949 |
| E | Together | Same generation | -0.134 | .949 |
| F | Alone | Intergenerational | 0.106 | .949 |
| F | Alone | Same generation | 1.548 | .341 |
| F | Together | Intergenerational | 2.679 | .052 |
| F | Together | Same generation | -0.191 | .949 |
| G | Alone | Intergenerational | -0.896 | .652 |
| G | Alone | Same generation | -0.235 | .949 |
| G | Together | Intergenerational | -3.298 | .020* |
| G | Together | Same generation | 0.054 | .957 |

*Note*. $p$-values were adjusted for multiple comparisons using the false discovery rate (FDR).

**Supplementary Table 10.** Likelihood ratio test results with Chi-squared tests for coverage, only real dyads.

| Factor | *df* | Δ Deviance ($\chi^2$) | $p$ |
|---|---|---|---|
| Brain state | 2 | 6.77 | .034* |
| Task | 1 | 0.77 | .381 |
| Group | 1 | 0.02 | .875 |
| Brain State × Task | 2 | 3.02 | .221 |
| Brain State × Group | 2 | 9.15 | .010* |
| Task × Group | 1 | 4.89 | .027* |
| Brain State × Task × Group | 2 | 1.04 | .594 |

*Note.* Deviance tests are likelihood ratio $\chi^2$-tests from a negative binomial GLM (log link; $\theta = 1.28$). The dependent variable is coverage in states B, D, and G.

**Supplementary Table 11.** Contrast analyses for coverage in states B, D and G, only real dyads.

| Contrast | State | $z$ | $p_{adj}$ |
|---|---|---|---|
| **Within-group task contrasts** | | | |
| Intergen: Alone vs. Together | B | -1.678 | .280 |
| Same gen: Alone vs. Together | B | -0.504 | .742 |
| Intergen: Alone vs. Together | D | -2.412 | .095 |
| Same gen: Alone vs. Together | D | 0.428 | .752 |
| Intergen: Alone vs. Together | G | 0.003 | .997 |
| Same gen: Alone vs. Together | G | 1.281 | .400 |
| **Within-task group contrasts** | | | |
| Alone: Intergen vs. Same gen | B | -1.057 | .523 |
| Together: Intergen vs. Same gen | B | 0.052 | .997 |
| Alone: Intergen vs. Same gen | D | -2.900 | .067 |
| Together: Intergen vs. Same gen | D | -0.498 | .742 |
| Alone: Intergen vs. Same gen | G | 0.574 | .742 |
| Together: Intergen vs. Same gen | G | 2.297 | .097 |

*Note. p*-values were adjusted for multiple comparisons using the false discovery rate (FDR).

**Supplementary Table 12.** Likelihood ratio test results with Chi-squared tests for duration, real and pseudo dyads.

| **Factor** | ***df*** | **Δ Deviance ($\chi^2$)** | ***p*** |
|---|---|---|---|
| Brain State | 6 | 1236.57 | < .001*** |
| Task | 1 | 7.37 | .007** |
| Dyad Type | 1 | 5.62 | .018* |
| Group | 1 | 0.24 | .622 |
| Session | 1 | 6.76 | .009** |
| Brain State × Task | 6 | 59.88 | < .001*** |
| Brain State × Dyad Type | 6 | 10.24 | .115 |
| Task × Dyad Type | 1 | 1.78 | .182 |
| Brain State × Group | 6 | 21.32 | .002** |
| Task × Group | 1 | 3.40 | .065 |
| Dyad Type × Group | 1 | 0.15 | .703 |
| Brain State × Session | 6 | 25.49 | < .001*** |
| Task × Session | 1 | 0.01 | .933 |
| Dyad Type × Session | 1 | 0.06 | .801 |
| Group × Session | 1 | 2.94 | .086 |
| Brain State × Task × Dyad Type | 6 | 5.14 | .526 |
| Brain State × Task × Group | 6 | 11.44 | .076 |
| Brain State × Dyad Type × Group | 6 | 31.08 | < .001*** |
| Task × Dyad Type × Group | 1 | 0.00 | .979 |
| Brain State × Task × Session | 6 | 15.26 | .018* |
| Brain State × Dyad Type × Session | 6 | 20.20 | .003** |
| Task × Dyad Type × Session | 1 | 0.01 | .930 |
| Brain State × Group × Session | 6 | 21.44 | .002** |
| Task × Group × Session | 1 | 1.12 | .291 |
| Dyad Type × Group × Session | 1 | 0.02 | .891 |
| Brain State × Task × Dyad Type × Group | 6 | 14.35 | .026* |
| Brain State × Task × Dyad Type × Session | 6 | 2.97 | .812 |
| Brain State × Task × Group × Session | 6 | 5.94 | .430 |
| Brain State × Dyad Type × Group × Session | 6 | 5.71 | .456 |
| Task × Dyad Type × Group × Session | 1 | 1.05 | .305 |
| Brain State × Task × Dyad Type × Group × Session | 6 | 6.99 | .321 |

*Note.* Deviance tests are likelihood ratio $\chi^2$-tests from a negative binomial GLM (log link; $\theta = 4.26$). The dependent variable is occurrence.

**Supplementary Table 13.** Contrast analyses for duration, real vs. pseudo dyads.

| Brain State | Task | Group | $z$ | $p_{adj}$ |
|---|---|---|---|---|
| A | Alone | Intergenerational | -0.671 | .827 |
| A | Alone | Same generation | 0.943 | .690 |
| A | Together | Intergenerational | -0.170 | .968 |
| A | Together | Same generation | 1.219 | .567 |
| B | Alone | Intergenerational | -0.126 | .968 |
| B | Alone | Same generation | -1.883 | .294 |
| B | Together | Intergenerational | -1.461 | .504 |
| B | Together | Same generation | 0.229 | .955 |
| C | Alone | Intergenerational | -0.735 | .809 |
| C | Alone | Same generation | -2.46 | .130 |
| C | Together | Intergenerational | 2.312 | .146 |
| C | Together | Same generation | -1.547 | .487 |
| D | Alone | Intergenerational | 1.149 | .585 |
| D | Alone | Same generation | -1.045 | .638 |
| D | Together | Intergenerational | 1.367 | .534 |
| D | Together | Same generation | -0.314 | .917 |
| E | Alone | Intergenerational | -2.471 | .130 |
| E | Alone | Same generation | -0.084 | .968 |
| E | Together | Intergenerational | 0.345 | .917 |
| E | Together | Same generation | 0.029 | .977 |
| F | Alone | Intergenerational | -1.294 | .547 |
| F | Alone | Same generation | -0.393 | .917 |
| F | Together | Intergenerational | 1.859 | .294 |
| F | Together | Same generation | -0.492 | .917 |
| G | Alone | Intergenerational | -0.897 | .690 |
| G | Alone | Same generation | -0.357 | .917 |
| G | Together | Intergenerational | -4.508 | < .001*** |
| G | Together | Same generation | 0.370 | .917 |

*Note*. *p*-values were adjusted for multiple comparisons using the false discovery rate (FDR).

**Supplementary Table 14.** Likelihood ratio test results with Chi-squared tests for duration, only real dyads.

| Factor | *df* | Δ Deviance ($\chi^2$) | *p* |
|---|---|---|---|
| Task | 1 | 0.23 | .630 |
| Group | 1 | 10.94 | <.001*** |
| Task × Group | 1 | 2.43 | .119 |

*Note.* Deviance tests are likelihood ratio $\chi^2$-tests from a negative binomial GLM (log link; $\theta = 2.98$). The dependent variable is duration in state G.

**Supplementary Table 15.** Contrast analyses for duration in state G, only real dyads.

| Contrast | State | $z$ | $p_{adj}$ |
|---|---|---|---|
| **Within-group task contrasts** | | | |
| Intergen: Alone vs. Together | G | -1.382 | .334 |
| Same gen: Alone vs. Together | G | 0.814 | .613 |
| **Within-task group contrasts** | | | |
| Alone: Intergen vs. Same gen | G | 0.658 | .613 |
| Together: Intergen vs. Same gen | G | 3.590 | .002** |

*Note. p*-values were adjusted for multiple comparisons using the false discovery rate (FDR).

**Supplementary Table 16.** Contrast analyses for slope of occurrence across sessions, real vs. pseudo dyads.

| Brain State | Task | Group | $z$ | $p_{adj}$ |
|---|---|---|---|---|
| A | Alone | Intergenerational | 0.148 | .995 |
| A | Alone | Same generation | 0.387 | .978 |
| A | Together | Intergenerational | 1.427 | .537 |
| A | Together | Same generation | 0.910 | .781 |
| B | Alone | Intergenerational | 0.084 | .995 |
| B | Alone | Same generation | 2.224 | .366 |
| B | Together | Intergenerational | 2.048 | .379 |
| B | Together | Same generation | -1.059 | .680 |
| C | Alone | Intergenerational | 0.595 | .909 |
| C | Alone | Same generation | -1.673 | .529 |
| C | Together | Intergenerational | -0.784 | .820 |
| C | Together | Same generation | -0.750 | .820 |
| D | Alone | Intergenerational | -1.055 | .680 |
| D | Alone | Same generation | -0.071 | .995 |
| D | Together | Intergenerational | 0.470 | .940 |
| D | Together | Same generation | 0.310 | .995 |
| E | Alone | Intergenerational | -1.448 | .537 |
| E | Alone | Same generation | -2.249 | .366 |
| E | Together | Intergenerational | -0.724 | .820 |
| E | Together | Same generation | -0.526 | .932 |
| F | Alone | Intergenerational | -0.007 | .995 |
| F | Alone | Same generation | 0.034 | .995 |
| F | Together | Intergenerational | -0.261 | .995 |
| F | Together | Same generation | 1.330 | .571 |
| G | Alone | Intergenerational | 0.123 | .995 |
| G | Alone | Same generation | -1.672 | .529 |
| G | Together | Intergenerational | -1.529 | .537 |
| G | Together | Same generation | 1.245 | .596 |

*Note. p*-values were adjusted for multiple comparisons using the false discovery rate (FDR).

**Supplementary Table 17.** Contrast analyses for slope of coverage across sessions, real vs. pseudo dyads.

| Brain State | Task | Group | $z$ | $p_{adj}$ |
|---|---|---|---|---|
| A | Alone | Intergenerational | -0.095 | .924 |
| A | Alone | Same generation | 0.869 | .611 |
| A | Together | Intergenerational | 0.644 | .633 |
| A | Together | Same generation | 1.080 | .611 |
| B | Alone | Intergenerational | 0.467 | .718 |
| B | Alone | Same generation | 1.416 | .611 |
| B | Together | Intergenerational | 0.733 | .611 |
| B | Together | Same generation | -0.860 | .611 |
| C | Alone | Intergenerational | 1.123 | .611 |
| C | Alone | Same generation | -0.989 | .611 |
| C | Together | Intergenerational | 0.756 | .611 |
| C | Together | Same generation | -1.571 | .611 |
| D | Alone | Intergenerational | 0.297 | .825 |
| D | Alone | Same generation | -0.968 | .611 |
| D | Together | Intergenerational | -0.591 | .647 |
| D | Together | Same generation | -0.984 | .611 |
| E | Alone | Intergenerational | -2.218 | .372 |
| E | Alone | Same generation | -2.646 | .228 |
| E | Together | Intergenerational | -1.580 | .611 |
| E | Together | Same generation | -1.157 | .611 |
| F | Alone | Intergenerational | 1.481 | .611 |
| F | Alone | Same generation | 1.284 | .611 |
| F | Together | Intergenerational | 0.706 | .611 |
| F | Together | Same generation | 1.398 | .611 |
| G | Alone | Intergenerational | -0.914 | .611 |
| G | Alone | Same generation | -0.241 | .840 |
| G | Together | Intergenerational | -0.744 | .611 |
| G | Together | Same generation | 0.719 | .611 |

*Note*. $p$-values were adjusted for multiple comparisons using the false discovery rate (FDR).

**Supplementary Table 18.** Contrast analyses for slope of duration across sessions, real vs. pseudo dyads.

| Brain State | Task | Group | $z$ | $p_{adj}$ |
|---|---|---|---|---|
| A | Alone | Intergenerational | -0.224 | .941 |
| A | Alone | Same generation | 0.935 | .624 |
| A | Together | Intergenerational | 0.296 | .941 |
| A | Together | Same generation | 0.325 | .941 |
| B | Alone | Intergenerational | 1.018 | .624 |
| B | Alone | Same generation | -0.813 | .624 |
| B | Together | Intergenerational | -1.491 | .624 |
| B | Together | Same generation | -0.074 | .941 |
| C | Alone | Intergenerational | 0.104 | .941 |
| C | Alone | Same generation | -0.662 | .712 |
| C | Together | Intergenerational | 0.891 | .624 |
| C | Together | Same generation | -1.036 | .624 |
| D | Alone | Intergenerational | 0.808 | .624 |
| D | Alone | Same generation | -1.158 | .624 |
| D | Together | Intergenerational | -1.275 | .624 |
| D | Together | Same generation | -1.062 | .624 |
| E | Alone | Intergenerational | -1.608 | .624 |
| E | Alone | Same generation | -1.703 | .624 |
| E | Together | Intergenerational | -1.212 | .624 |
| E | Together | Same generation | 0.228 | .941 |
| F | Alone | Intergenerational | 3.116 | .051 |
| F | Alone | Same generation | 1.128 | .624 |
| F | Together | Intergenerational | 1.113 | .624 |
| F | Together | Same generation | 0.896 | .624 |
| G | Alone | Intergenerational | -1.146 | .624 |
| G | Alone | Same generation | 0.800 | .624 |
| G | Together | Intergenerational | -0.193 | .941 |
| G | Together | Same generation | 0.093 | .941 |

*Note*. $p$-values were adjusted for multiple comparisons using the false discovery rate (FDR).

**Supplementary Table 19.** Model comparisons of Poisson and negative binomial models in the three-cluster solution (SR1).

| Metric | Model | Poisson model | Negative binomial model | |
|---|---|---|---|---|
| | | **AIC** | **θ** | **AIC** |
| Occurrence | First step | 56569.8 | – | 56571.86 |
| Occurrence | Second step | 1280.9 | – | 1282.9 |
| Coverage | First step | 427824.7 | 2.69 | 129576.5 |
| Coverage | Second step | 11250.3 | 1.85 | 2922.9 |
| Duration | First step | 123934.5 | 3.67 | 84967.3 |

*Note.* First-step models refer to generalized linear models to test differences between real and pseudo dyads. Second-step models refer to models constructed only on data from real dyads and those states that differed between pseudo and real dyads in at least one condition in first-step models. The dispersion parameter theta could not be estimated for the negative binomial models of occurrence. AIC = Akaike Information Criterion.

**Supplementary Table 20.** Likelihood ratio test results with Chi-squared tests for occurrence in the three-cluster solution (SR1), real and pseudo dyads.

| Factor | *df* | Δ Deviance ($\chi^2$) | *p* |
|---|---|---|---|
| Brain State | 6 | 999.45 | < .001*** |
| Task | 1 | 1.43 | .232 |
| Dyad Type | 1 | 0.02 | .895 |
| Group | 1 | 22.27 | < .001*** |
| Session | 1 | 6.58 | .010* |
| Brain State × Task | 6 | 17.05 | < .001*** |
| Brain State × Dyad Type | 6 | 7.14 | .028* |
| Task × Dyad Type | 1 | 4.88 | .027* |
| Brain State × Group | 6 | 4.51 | .105 |
| Task × Group | 1 | 2.21 | .137 |
| Dyad Type × Group | 1 | 1.99 | .158 |
| Brain State × Session | 6 | 1.34 | .511 |
| Task × Session | 1 | 10.74 | .001** |
| Dyad Type × Session | 1 | 0.00 | .973 |
| Group × Session | 1 | 1.78 | .183 |
| Brain State × Task × Dyad Type | 6 | 0.96 | .617 |
| Brain State × Task × Group | 6 | 0.28 | .869 |
| Brain State × Dyad Type × Group | 6 | 7.99 | .018* |
| Task × Dyad Type × Group | 1 | 0.06 | .800 |
| Brain State × Task × Session | 6 | 2.60 | .273 |
| Brain State × Dyad Type × Session | 6 | 1.73 | .421 |
| Task × Dyad Type × Session | 1 | 0.08 | .774 |
| Brain State × Group × Session | 6 | 2.56 | .279 |
| Task × Group × Session | 1 | 1.63 | .202 |
| Dyad Type × Group × Session | 1 | 2.02 | .155 |
| Brain State × Task × Dyad Type × Group | 6 | 1.00 | .608 |
| Brain State × Task × Dyad Type × Session | 6 | 0.18 | .915 |
| Brain State × Task × Group × Session | 6 | 1.57 | .457 |
| Brain State × Dyad Type × Group × Session | 6 | 0.24 | .885 |
| Task × Dyad Type × Group × Session | 1 | 0.54 | .463 |
| Brain State × Task × Dyad Type × Group × Session | 6 | 1.67 | .434 |

*Note.* Deviance tests are likelihood ratio $\chi^2$-tests from a Poisson GLM (log link). The dependent variable is occurrence.

**Supplementary Table 21.** Contrast analyses for occurrence in the three-cluster solution (SR1), real vs. pseudo dyads.

| **Brain State** | **Task** | **Group** | $z$ | $p_{adj}$ |
|---|---|---|---|---|
| I | Alone | Intergenerational | 0.335 | .769 |
| I | Alone | Same generation | 1.691 | .272 |
| I | Together | Intergenerational | -2.055 | .160 |
| I | Together | Same generation | 0.708 | .718 |
| II | Alone | Intergenerational | 0.294 | .769 |
| II | Alone | Same generation | 0.526 | .732 |
| II | Together | Intergenerational | 1.297 | .467 |
| II | Together | Same generation | -1.107 | .529 |
| III | Alone | Intergenerational | -1.018 | .529 |
| III | Alone | Same generation | 0.510 | .732 |
| III | Together | Intergenerational | -3.576 | .004** |
| III | Together | Same generation | -2.355 | .111 |

*Note*. *p*-values were adjusted for multiple comparisons using the false discovery rate (FDR).

**Supplementary Table 22.** Likelihood ratio test results with Chi-squared tests for occurrence in the three-cluster solution (SR1), only real dyads.

| **Factor** | ***df*** | **Δ Deviance ($\chi^2$)** | ***p*** |
|---|---|---|---|
| Task | 1 | 3.30 | .069 |
| Group | 1 | 0.18 | .668 |
| Task × Group | 1 | 0.18 | .668 |

*Note.* Deviance tests are likelihood ratio $\chi^2$-tests from a Poisson GLM (log link). The dependent variable is occurrence in state III.

**Supplementary Table 23.** Likelihood ratio test results with Chi-squared tests for coverage in the three-cluster solution (SR1), real and pseudo dyads.

| **Factor** | ***df*** | **Δ Deviance ($\chi^2$)** | ***p*** |
|---|---|---|---|
| Brain State | 6 | 1212.78 | < .001*** |
| Task | 1 | 2.68 | .102 |
| Dyad Type | 1 | 0.29 | .588 |
| Group | 1 | 0.28 | .594 |
| Session | 1 | 0.04 | .835 |
| Brain State × Task | 6 | 68.10 | < .001*** |
| Brain State × Dyad Type | 6 | 10.39 | .006** |
| Task × Dyad Type | 1 | 0.03 | .862 |
| Brain State × Group | 6 | 24.32 | < .001*** |
| Task × Group | 1 | 0.13 | .723 |
| Dyad Type × Group | 1 | 0.00 | .956 |
| Brain State × Session | 6 | 5.48 | .065 |
| Task × Session | 1 | 0.01 | .938 |
| Dyad Type × Session | 1 | 0.27 | .602 |
| Group × Session | 1 | 0.00 | .947 |
| Brain State × Task × Dyad Type | 6 | 0.08 | .959 |
| Brain State × Task × Group | 6 | 4.49 | .106 |
| Brain State × Dyad Type × Group | 6 | 19.11 | < .001*** |
| Task × Dyad Type × Group | 1 | 0.05 | .817 |
| Brain State × Task × Session | 6 | 1.32 | .517 |
| Brain State × Dyad Type × Session | 6 | 15.74 | < .001*** |
| Task × Dyad Type × Session | 1 | 0.12 | .727 |
| Brain State × Group × Session | 6 | 2.98 | .225 |
| Task × Group × Session | 1 | 0.00 | .996 |
| Dyad Type × Group × Session | 1 | 0.01 | .908 |
| Brain State × Task × Dyad Type × Group | 6 | 0.40 | .818 |
| Brain State × Task × Dyad Type × Session | 6 | 1.60 | .450 |
| Brain State × Task × Group × Session | 6 | 0.02 | .988 |
| Brain State × Dyad Type × Group × Session | 6 | 2.68 | .262 |
| Task × Dyad Type × Group × Session | 1 | 0.01 | .929 |
| Brain State × Task × Dyad Type × Group × Session | 6 | 2.26 | .324 |

*Note.* Deviance tests are likelihood ratio $\chi^2$-tests from a negative binomial GLM (log link; $\theta = 2.69$). The dependent variable is coverage.

**Supplementary Table 24.** Contrast analyses for coverage in the three-cluster solution (SR1), real vs. pseudo dyads.

| Brain State | Task | Group | *z* | $p_{adj}$ |
|---|---|---|---|---|
| I | Alone | Intergenerational | 0.418 | .713 |
| I | Alone | Same generation | 1.885 | .120 |
| I | Together | Intergenerational | -0.367 | .713 |
| I | Together | Same generation | 2.039 | .120 |
| II | Alone | Intergenerational | 1.358 | .299 |
| II | Alone | Same generation | -1.055 | .437 |
| II | Together | Intergenerational | 2.886 | .047* |
| II | Together | Same generation | -0.869 | .474 |
| III | Alone | Intergenerational | -2.118 | .120 |
| III | Alone | Same generation | -0.851 | .474 |
| III | Together | Intergenerational | -2.388 | .102 |
| III | Together | Same generation | -1.883 | .120 |

*Note*. *p*-values were adjusted for multiple comparisons using the false discovery rate (FDR).

**Supplementary Table 25.** Likelihood ratio test results with Chi-squared tests for coverage, only real dyads.

| Factor | *df* | Δ Deviance ($\chi^2$) | *p* |
|---|---|---|---|
| Task | 1 | 0.42 | .515 |
| Group | 1 | 6.12 | .013* |
| Task × Group | 1 | 0.23 | .628 |

*Note.* Deviance tests are likelihood ratio $\chi^2$-tests from a negative binomial GLM (log link; θ = 1.85). The dependent variable is coverage in state II.

**Supplementary Table 26.** Contrast analyses for coverage in state II in the three-cluster solution (SR1), only real dyads.

| Contrast | State | *z* | $p_{adj}$ |
|---|---|---|---|
| **Within-group task contrasts** | | | |
| Intergen: Alone vs. Together | II | 0.054 | .957 |
| Same gen: Alone vs. Together | II | 0.780 | .653 |
| **Within-task group contrasts** | | | |
| Alone: Intergen vs. Same gen | II | -1.835 | .162 |
| Together: Intergen vs. Same gen | II | -1.746 | .162 |

*Note. p*-values were adjusted for multiple comparisons using the false discovery rate (FDR).

**Supplementary Table 27.** Likelihood ratio test results with Chi-squared tests for duration in the three-cluster solution (SR1), real and pseudo dyads.

| **Factor** | ***df*** | **Δ Deviance ($\chi^2$)** | ***p*** |
|---|---|---|---|
| Brain State | 6 | 260.41 | < .001*** |
| Task | 1 | 4.29 | .038* |
| Dyad Type | 1 | 0.04 | .839 |
| Group | 1 | 5.59 | .018* |
| Session | 1 | 8.45 | .004** |
| Brain State × Task | 6 | 62.13 | < .001*** |
| Brain State × Dyad Type | 6 | 2.87 | .238 |
| Task × Dyad Type | 1 | 3.82 | .051 |
| Brain State × Group | 6 | 31.66 | < .001*** |
| Task × Group | 1 | 2.31 | .128 |
| Dyad Type × Group | 1 | 0.03 | .859 |
| Brain State × Session | 6 | 14.01 | .001** |
| Task × Session | 1 | 6.20 | .013* |
| Dyad Type × Session | 1 | 0.64 | .424 |
| Group × Session | 1 | 1.00 | .318 |
| Brain State × Task × Dyad Type | 6 | 0.05 | .976 |
| Brain State × Task × Group | 6 | 6.44 | .040* |
| Brain State × Dyad Type × Group | 6 | 9.98 | .007** |
| Task × Dyad Type × Group | 1 | 0.01 | .920 |
| Brain State × Task × Session | 6 | 0.54 | .763 |
| Brain State × Dyad Type × Session | 6 | 12.54 | .002** |
| Task × Dyad Type × Session | 1 | 0.01 | .921 |
| Brain State × Group × Session | 6 | 1.86 | .395 |
| Task × Group × Session | 1 | 3.47 | .062 |
| Dyad Type × Group × Session | 1 | 1.81 | .178 |
| Brain State × Task × Dyad Type × Group | 6 | 3.50 | .174 |
| Brain State × Task × Dyad Type × Session | 6 | 4.07 | .130 |
| Brain State × Task × Group × Session | 6 | 0.13 | .936 |
| Brain State × Dyad Type × Group × Session | 6 | 1.16 | .559 |
| Task × Dyad Type × Group × Session | 1 | 0.17 | .683 |
| Brain State × Task × Dyad Type × Group × Session | 6 | 0.49 | .783 |

*Note.* Deviance tests are likelihood ratio $\chi^2$-tests from a negative binomial GLM (log link; $\theta = 3.67$). The dependent variable is occurrence.

**Supplementary Table 28.** Contrast analyses for duration in the three-cluster solution (SR1), real vs. pseudo dyads.

| Brain State | Task | Group | $z$ | $p_{adj}$ |
|---|---|---|---|---|
| I | Alone | Intergenerational | 0.050 | .960 |
| I | Alone | Same generation | 1.277 | .484 |
| I | Together | Intergenerational | 1.040 | .511 |
| I | Together | Same generation | 2.472 | .111 |
| II | Alone | Intergenerational | 1.095 | .511 |
| II | Alone | Same generation | -2.350 | .111 |
| II | Together | Intergenerational | 2.085 | .111 |
| II | Together | Same generation | 0.574 | .755 |
| III | Alone | Intergenerational | -2.179 | .111 |
| III | Alone | Same generation | -0.615 | .755 |
| III | Together | Intergenerational | -0.465 | .768 |
| III | Together | Same generation | -0.380 | .768 |

*Note*. *p*-values were adjusted for multiple comparisons using the false discovery rate (FDR).

**Supplementary Table 29.** Contrast analyses for slope of occurrence across sessions in the three-cluster solution (SR1), real vs. pseudo dyads.

| Brain State | Task | Group | $z$ | $p_{adj}$ |
|---|---|---|---|---|
| I | Alone | Intergenerational | 0.573 | .927 |
| I | Alone | Same generation | 0.825 | .927 |
| I | Together | Intergenerational | 1.087 | .927 |
| I | Together | Same generation | 0.284 | .927 |
| II | Alone | Intergenerational | 1.430 | .916 |
| II | Alone | Same generation | -0.889 | .927 |
| II | Together | Intergenerational | 0.151 | .927 |
| II | Together | Same generation | 0.092 | .927 |
| III | Alone | Intergenerational | 0.092 | .927 |
| III | Alone | Same generation | -1.472 | .916 |
| III | Together | Intergenerational | 0.325 | .927 |
| III | Together | Same generation | -0.392 | .927 |

*Note*. *p*-values were adjusted for multiple comparisons using the false discovery rate (FDR).

**Supplementary Table 30.** Contrast analyses for slope of coverage across sessions in the three-cluster solution (SR1), real vs. pseudo dyads.

| Brain State | Task | Group | $z$ | $p_{adj}$ |
|---|---|---|---|---|
| I | Alone | Intergenerational | 0.245 | .879 |
| I | Alone | Same generation | 1.481 | .334 |
| I | Together | Intergenerational | 1.079 | .421 |
| I | Together | Same generation | 1.158 | .421 |
| II | Alone | Intergenerational | 1.916 | .332 |
| II | Alone | Same generation | -0.512 | .730 |
| II | Together | Intergenerational | -0.085 | .932 |
| II | Together | Same generation | 0.729 | .622 |
| III | Alone | Intergenerational | -2.158 | .332 |
| III | Alone | Same generation | -1.353 | .352 |
| III | Together | Intergenerational | -1.690 | .334 |
| III | Together | Same generation | -1.479 | .334 |

*Note*. $p$-values were adjusted for multiple comparisons using the false discovery rate (FDR).

**Supplementary Table 31.** Contrast analyses for slope of duration across sessions in the three-cluster solution (SR1), real vs. pseudo dyads.

| Brain State | Task | Group | $z$ | $p_{adj}$ |
|---|---|---|---|---|
| I | Alone | Intergenerational | 0.383 | .766 |
| I | Alone | Same generation | 0.915 | .550 |
| I | Together | Intergenerational | 0.191 | .848 |
| I | Together | Same generation | 0.922 | .550 |
| II | Alone | Intergenerational | 0.742 | .550 |
| II | Alone | Same generation | 0.990 | .550 |
| II | Together | Intergenerational | -0.786 | .550 |
| II | Together | Same generation | -0.771 | .550 |
| III | Alone | Intergenerational | -2.796 | .062 |
| III | Alone | Same generation | -1.072 | .550 |
| III | Together | Intergenerational | -1.994 | .277 |
| III | Together | Same generation | -1.249 | .550 |

*Note*. $p$-values were adjusted for multiple comparisons using the false discovery rate (FDR).

**Supplementary Table 32.** Likelihood ratio test results with Chi-squared tests for coverage of shared brain states in the three-cluster solution (SR2), real and pseudo dyads.

| **Factor** | ***df*** | **Δ Deviance ($\chi^2$)** | ***p*** |
|---|---|---|---|
| Brain State | 6 | 4633.92 | < .001*** |
| Task | 1 | 8372.50 | < .001*** |
| Dyad Type | 1 | 0.01 | .931 |
| Group | 1 | 0.35 | .556 |
| Session | 1 | 0.81 | .367 |
| Brain State × Task | 6 | 28.42 | < .001*** |
| Brain State × Dyad Type | 6 | 0.17 | .682 |
| Task × Dyad Type | 1 | 0.18 | .669 |
| Brain State × Group | 6 | 6.13 | .013* |
| Task × Group | 1 | 0.001 | .921 |
| Dyad Type × Group | 1 | 0.52 | .469 |
| Brain State × Session | 6 | 14.40 | < .001*** |
| Task × Session | 1 | 0.05 | .831 |
| Dyad Type × Session | 1 | 0.03 | .871 |
| Group × Session | 1 | 0.01 | .943 |
| Brain State × Task × Dyad Type | 6 | 3.19 | .074 |
| Brain State × Task × Group | 6 | 0.05 | .825 |
| Brain State × Dyad Type × Group | 6 | 9.43 | .002** |
| Task × Dyad Type × Group | 1 | 0.05 | .822 |
| Brain State × Task × Session | 6 | 0.31 | .578 |
| Brain State × Dyad Type × Session | 6 | 0.57 | .452 |
| Task × Dyad Type × Session | 1 | 0.07 | .792 |
| Brain State × Group × Session | 6 | 0.02 | .875 |
| Task × Group × Session | 1 | 0.00 | .987 |
| Dyad Type × Group × Session | 1 | 0.08 | .776 |
| Brain State × Task × Dyad Type × Group | 6 | 1.11 | .292 |
| Brain State × Task × Dyad Type × Session | 6 | 1.10 | .294 |
| Brain State × Task × Group × Session | 6 | 0.00 | .956 |
| Brain State × Dyad Type × Group × Session | 6 | 0.94 | .332 |
| Task × Dyad Type × Group × Session | 1 | 0.00 | .968 |
| Brain State × Task × Dyad Type × Group × Session | 6 | 0.05 | .821 |

*Note.* Deviance tests are likelihood ratio $\chi^2$-tests from a negative binomial GLM (log link; θ = 16.17). The dependent variable is occurrence.

**Supplementary Materials:** [redacted for peer review]. *Two-brain states during collaborative drawing reflect leader–follower dynamics in intergenerational dyads.*

**Supplementary Table 33.** Contrast analyses for coverage of shared brain states in the three-cluster solution (SR2), real vs. pseudo dyads.

| Brain State | Task | Group | $z$ | $p_{adj}$ |
|---|---|---|---|---|
| shared | Alone | Intergenerational | 1.623 | .279 |
| shared | Alone | Same generation | -0.458 | .806 |
| shared | Together | Intergenerational | 0.246 | .806 |
| shared | Together | Same generation | -0.948 | .649 |
| non-shared | Alone | Intergenerational | -2.547 | .087 |
| non-shared | Alone | Same generation | 0.832 | .649 |
| non-shared | Together | Intergenerational | -0.369 | .806 |
| non-shared | Together | Same generation | 1.670 | .279 |

*Note*. *p*-values were adjusted for multiple comparisons using the false discovery rate (FDR).

**Supplementary Table 34.** Likelihood ratio test results with Chi-squared tests for coverage of shared brain states in the eight-cluster solution (SR2), real and pseudo dyads.

| **Factor** | ***df*** | **Δ Deviance ($\chi^2$)** | ***p*** |
|---|---|---|---|
| Brain State | 6 | 38176.55 | < .001*** |
| Task | 1 | 5953.91 | < .001*** |
| Dyad Type | 1 | 1.21 | .271 |
| Group | 1 | 1.79 | .181 |
| Session | 1 | 2.58 | .108 |
| Brain State × Task | 6 | 2.09 | .148 |
| Brain State × Dyad Type | 6 | 2.83 | .092 |
| Task × Dyad Type | 1 | 0.12 | .733 |
| Brain State × Group | 6 | 4.21 | .040* |
| Task × Group | 1 | 0.14 | .705 |
| Dyad Type × Group | 1 | 0.22 | .636 |
| Brain State × Session | 6 | 6.45 | .011* |
| Task × Session | 1 | 0.41 | .523 |
| Dyad Type × Session | 1 | 0.23 | .632 |
| Group × Session | 1 | 0.43 | .513 |
| Brain State × Task × Dyad Type | 6 | 0.30 | .583 |
| Brain State × Task × Group | 6 | 0.36 | .549 |
| Brain State × Dyad Type × Group | 6 | 0.56 | .455 |
| Task × Dyad Type × Group | 1 | 0.20 | .655 |
| Brain State × Task × Session | 6 | 0.98 | .321 |
| Brain State × Dyad Type × Session | 6 | 0.47 | .492 |
| Task × Dyad Type × Session | 1 | 1.01 | .315 |
| Brain State × Group × Session | 6 | 1.04 | .307 |
| Task × Group × Session | 1 | 0.21 | .647 |
| Dyad Type × Group × Session | 1 | 0.03 | .861 |
| Brain State × Task × Dyad Type × Group | 6 | 0.50 | .479 |
| Brain State × Task × Dyad Type × Session | 6 | 2.56 | .110 |
| Brain State × Task × Group × Session | 6 | 0.54 | .464 |
| Brain State × Dyad Type × Group × Session | 6 | 0.08 | .776 |
| Task × Dyad Type × Group × Session | 1 | 0.26 | .609 |
| Brain State × Task × Dyad Type × Group × Session | 6 | 0.63 | .428 |

*Note.* Deviance tests are likelihood ratio $\chi^2$-tests from a negative binomial GLM (log link; $\theta = 12.09$). The dependent variable is occurrence.

**Supplementary Table 35.** Contrast analyses for coverage of shared brain states in the eight-cluster solution (SR2), real vs. pseudo dyads.

| **Brain State** | **Task** | **Group** | $z$ | $p_{adj}$ |
|---|---|---|---|---|
| shared | Alone | Intergenerational | -0.093 | .926 |
| shared | Alone | Same generation | 0.241 | .926 |
| shared | Together | Intergenerational | 0.223 | .926 |
| shared | Together | Same generation | 0.218 | .926 |
| non-shared | Alone | Intergenerational | 0.605 | .926 |
| non-shared | Alone | Same generation | -1.358 | .572 |
| non-shared | Together | Intergenerational | -1.241 | .572 |
| non-shared | Together | Same generation | -1.245 | .572 |

*Note*. *p*-values were adjusted for multiple comparisons using the false discovery rate (FDR).

**Supplementary Materials:** [redacted for peer review]. *Two-brain states during collaborative drawing reflect leader–follower dynamics in intergenerational dyads.*

**Supplementary Table 36.** Regression for percentage of turn taking during collaboration in real dyads.

| **Factor** | ***b*** | ***SE*** | ***t*** | ***p*** |
|---|---|---|---|---|
| Intercept | 4.224 | 0.509 | 8.294 | < .001*** |
| Duration of state G | -0.069 | 0.023 | -3.051 | .003** |
| Group | -1.602 | 0.492 | -3.254 | .001** |
| Session | -0.111 | 0.093 | -1.201 | .232 |
| Duration of state G*Group | 0.052 | 0.052 | -1.008 | .315 |

*Note*. $F(4, 149) = 5.797$, $p \leq .001$. $R^2 = .11$.

**Supplementary Table 37.** Regression for duration of state G during collaboration in real, intergenerational dyads.

| **Factor** | ***b*** | ***SE*** | ***t*** | ***p*** |
|---|---|---|---|---|
| Intercept | 36.300 | 15.170 | 2.393 | .019* |
| Age difference | -0.406 | 0.267 | -1.519 | .133 |
| Session | -1.092 | 0.850 | -1.285 | .203 |

*Note*. $F(2, 71) = 1.617$, $p = .206$. $R^2 = .02$.

**Supplementary Table 38.** Regression for duration of state G during collaboration in real, intergenerational dyads.

| **Factor** | ***b*** | ***SE*** | ***t*** | ***p*** |
|---|---|---|---|---|
| Intercept | 53.068 | 26.304 | 2.018 | .047* |
| Age of older person | -0.501 | 0.333 | -1.502 | .138 |
| Session | -1.023 | 0.841 | -1.216 | .228 |

*Note*. $F(2, 71) = 1.591$, $p = .211$. $R^2 = .02$.

**Supplementary Table 39.** Regression for extent of turn-taking during collaboration in real, intergenerational dyads.

| Factor | *b* | *SE* | *t* | *p* |
|---|---|---|---|---|
| Intercept | 0.070 | 2.569 | 0.027 | .978 |
| Age difference | 0.064 | 0.045 | 1.405 | .165 |
| Session | -0.077 | 0.144 | -0.532 | .596 |

*Note*. $F(2, 71) = 1.371$, $p = .261$. $R^2 = .07$.

**Supplementary Table 40.** Regression for extent of turn-taking during collaboration in real, intergenerational dyads.

| Factor | *b* | *SE* | *t* | *p* |
|---|---|---|---|---|
| Intercept | -8.838 | 4.262 | -2.074 | .042* |
| Age of older person | 0.159 | 0.054 | 2.940 | .004** |
| Session | -0.051 | 0.136 | -0.375 | .709 |

*Note*. $F(2, 71) = 4.743$, $p = .012$. $R^2 = .09$.

**Supplementary Materials:** [redacted for peer review]. *Two-brain states during collaborative drawing reflect leader–follower dynamics in intergenerational dyads.*

**Supplementary Results**

**SR1 Three-cluster solution.**

We conducted the analyses of the three-cluster solution exactly analogously to the analyses of the seven-cluster solution. First, we identified the two-brain states that emerge during creative-art making within this sample. We set qualitative thresholds based on the distributions of connectivity values to describe the connectivity: high >= 0.3, medium >= 0.25, low >= 0.2, very low < 0.2. The three two-brain states are labelled I, II, and III. State I is characterised by low connectivity between brains and medium connectivity within brains. States II and III are characterised by medium-to-high connectivity between brains and high within-brain connectivity in only one dyad member (I for the first and III for the second dyad member).

For each brain-state and metric (occurrence, coverage, duration) and brain-state (A–G) we first assessed the extent to which the metric differs between real and pseudo dyads. Next, using planned contrasts, we evaluated the extent to which dyadic brain states differ between collaborative and independent drawing, intergenerational and same-generation dyads, and the interaction between these contexts. As for the seven-cluster solution, we tested for each analysis whether a Poisson model would be appropriate or if overdispersion necessitates an approached based on the negative binomial distribution. Model comparisons are reported in Supplementary Table 19.

**SR1.1 Occurrence (number of discrete entries into a brain state)**

**Real vs. pseudo dyads.** We observed differences between real and pseudo dyads for state III (Intergenerational group, drawing together: $z$ = -3.58, $p_{adj}$ = .004). States I and II did not differ significantly between real and pseudo dyads for any drawing condition or group (Supplementary Tables 20 and 21).

**State III.** Likelihood ratio tests showed no main or interaction effects ($p$s > .05; Supplementary Table 22). Therefore, we did not conduct any further analyses.

**SR1.2 Coverage (total number of windows in which a brain state is active)**

**Real vs. pseudo dyads.** We observed a difference between real and pseudo dyads for state II (Intergenerational group, drawing together: $z$ = -2898, $p_{adj}$ = .047). States I and III did not differ significantly between real and pseudo dyads for any drawing condition or group (Supplementary Tables 23 and 24).

**State II.** Likelihood ratio tests showed a main effect of group ($p$ < .05; Supplementary Table 25). Contrast analyses revealed no statistically significant differences in any states between tasks or groups after FDR correction (Supplementary Table 26).

**SR1.3 Duration (mean number of consecutive windows in which a brain state is active)**

**Real vs. pseudo dyads.** We observed no differences between real and pseudo dyads (Supplementary Tables 27 and 28), therefore we did not conduct any further analyses.

#### SR1.4 Longitudinal changes in two-brain states

We examined the differences in the slopes of occurrence, coverage, and duration of the two-brain states across sessions between real dyads and pseudo dyads. There were no significant differences in occurrence, duration or coverage after FDR correction (Supplementary Tables 29–31), therefore, we did not conduct any further analyses.

### SR2 Additional pre-registered hypotheses: Shared brain state analysis.

We additionally pre-registered an analysis of shared brain states analogous to (Li et al., 2025). In this pipeline, we clustered single-brain data and examined the proportion of time covered by windows with shared brain states (i.e., both members of a dyad show the same state) versus non-shared brain states (i.e., members of a dyad show different states). We again selected the best combination of hyperparameters using CHI and DBI; CHI indicated a three-cluster solution (RBF kernel, $\lambda = 0.1$), DBI indicated an eight-cluster solution (RBF kernel, $\lambda = 0.1$). Results for both solutions are reported below.

To establish whether each brain state was the result of true dyadic interactions or could also be observed when the signals of who did not interact were aligned, we contrasted coverage of shared brain states between real and pseudo dyads. The formula was the following: *coverage ~ state * drawingTask * dyadType * group * session,* where drawingTask refers to drawing together or alone, dyadType refers to real and pseudo dyads, group refers to same generation and intergeneration groups, and state was a binary variable indicating shared vs. non-shared brain states. Next, we planned to examine the extent to which the coverage of shared brain states differs between collaborative and independent drawing, intergenerational and same-generation dyads, and the interaction between these contexts. As for the two-brain analyses, we tested for each analysis whether a Poisson model would be appropriate or if overdispersion necessitates an approached based on the negative binomial distribution.

**Three-cluster solution.** Model fit was better for a negative binomial model (AIC = 49660.8) than for a Poisson model (AIC = 85461.8). We observed no differences between real and pseudo dyads, although there was a trend towards higher coverage for non-shared states in intergenerational dyads for drawing alone (Supplementary Tables 32 and 33). Therefore, we did not conduct any further analyses.

**Eight-cluster solution.** Model fit was better for a negative binomial model (AIC = 47589.3) than for a Poisson model (AIC = 61138.4). We observed no differences between real and pseudo dyads (Supplementary Tables 34 and 35). Therefore, we did not conduct any further analyses.

**Supplementary Materials:** [redacted for peer review]. *Two-brain states during collaborative drawing reflect leader–follower dynamics in intergenerational dyads.*

### References

Goes, J., Lerman, G., & Nadler, B. (2020). Robust sparse covariance estimation by thresholding Tyler's M-estimator. *The Annals of Statistics*, *48*(1). https://doi.org/10.1214/18-AOS1793

Ledoit, O., & Wolf, M. (2004). A well-conditioned estimator for large-dimensional covariance matrices. *Journal of Multivariate Analysis*, *88*(2), 365–411. https://doi.org/10.1016/S0047-259X(03)00096-4

Li, Q., Zimmermann, M., & Konvalinka, I. (2025). Two-brain microstates: A novel hyperscanning-EEG method for quantifying task-driven inter-brain asymmetry. *Journal of Neuroscience Methods*, *416*, 110355. https://doi.org/10.1016/j.jneumeth.2024.110355

Rocha, H. (2009). On the selection of the most adequate radial basis function. *Applied Mathematical Modelling*, *33*(3), 1573–1583. https://doi.org/10.1016/j.apm.2008.02.008

Ver Hoef, J. M., & Boveng, P. L. (2007). Quasi-Poisson vs. negative binomial regression: How should we model overdispersed count data? *Ecology*, *88*(11), 2766–2772. https://doi.org/10.1890/07-0043.1
